# Heterogeneity during induction of resistance delays *Pseudomonas aeruginosa* recovery from antibiotic exposures

**DOI:** 10.1101/2024.09.20.614194

**Authors:** David Ritz, Rachel Martin, Mabel Bush, JoAnne Villagrana, Yijie Deng, Daniel Schultz

## Abstract

Typical antibiotic susceptibility testing (AST) of microbial samples is performed in homogeneous cultures in batch environments, which does not account for the highly heterogeneous and dynamic nature of antibiotic responses. The most common mutation found in *P. aeruginosa* lineages evolved during chronic infections in the human lung, a loss of function of repressor MexZ, increases basal levels of multidrug efflux MexXY, but does not increase resistance by traditional minimal inhibitory concentration (MIC) assays. Here, we use single cell microfluidics to show that *P. aeruginosa* response to aminoglycosides is highly heterogeneous, with only a subpopulation of cells surviving exposure. In contrast, strains carrying *mexZ* mutations bypass the lengthy process of MexXY activation, increasing survival to sudden drug exposures and conferring a fitness advantage in fluctuating environments. Building on the data we present here, we propose a simple “Response Dynamics” assay to quantify the rate of population-level recovery to drug exposures across strains. We used this assay to profile a representative panel of 49 *P. aeruginosa* strains from diverse environments, showing that the presence of *mexZ* mutations correlates with faster population recovery from exposures to aminoglycosides, and thus confers an advantage to cells exposed to a sudden, large dose of antibiotic. We propose that the Response Dynamics assay can be used alongside MIC assays for profiling of antibiotic sensitivity to better predict clinical outcomes from *in vitro* sensitivity/resistance profiles.

**Significance:** Common mutations affecting the regulation of antibiotic resistance in bacterial pathogens often do not increase resistance by traditional measures. However, antibiotic resistance is typically measured in stable cultures, without accounting for fluctuations in drug concentration. Here, we show that *P. aeruginosa* response to aminoglycosides is highly heterogeneous, and that the most common mutation found in clinical isolates improves resistance by increasing single-cell survival to drug exposures. Therefore, the success of antibiotic treatments depends on the dynamics of drug delivery and the heterogeneous activation of microbial responses. We then develop an assay to measure resistance in a dynamic context, which captures this overlooked aspect of antibiotic resistance and can be used alongside traditional measures in the profiling of clinical isolates.

## Introduction

The human host presents a complex and dynamic environment for the evolution of opportunistic pathogens. While establishing infections, such microbes encounter various challenges, including antibiotic treatments wherein they are periodically exposed to large drug doses at concentrations much higher than typical exposures in nature^1^. Opportunistic pathogens are usually equipped with a variety of inducible mechanisms of antibiotic resistance, but sudden exposures require rapid responses while gene expression is still possible^2,3^. Consequently, not all cells are able to successfully mount a response to rapid, large-dose antibiotic challenges, which often leads to the co-occurrence of live and dead (or arrested) cells^4^. This heterogeneity influences treatment outcomes and complicates the utility of *in vitro* antibiotic susceptibility testing (AST) of pathogens as predictors of clinical outcomes^5–7^. Currently, susceptibility profiling of pathogens in the lab, while effective in many circumstances, typically only estimates the minimum inhibitory concentration (MIC) of antibiotics in bulk bacterial cultures. Recently, population analysis profiling (PAP)^8^ has been used to quantify phenotypic heterogeneity during antibiotic treatments (heteroresistance), but it still does not account for the noisy dynamics of drug responses. We hypothesize that accounting for such dynamics can increase the effectiveness of *in vitro* assays in predicting treatment outcomes of infections in the host.

*Pseudomonas aeruginosa*, an opportunistic bacterial pathogen, is a leading cause of infections in human hosts. It is particularly associated with cystic fibrosis (CF), a genetic disease that causes mucus accumulation in the lung, which becomes prone to colonization by bacteria, resulting in loss of lung function^9,10^. *P. aeruginosa* infections are typically acquired from unique environmental clone types naïve to the human host^11^, but lineages evolved in chronic infections eventually diversify into heterogeneous populations with remarkably altered phenotypes, including reduced virulence, transition to biofilm growth and reduced sensitivity to antibiotics^12–15^. Antibiotics, particularly aminoglycosides, have been routinely used to treat *P. aeruginosa* infections^16^. Although early infections can sometimes be cleared^17^, chronic infections are difficult to eradicate^18^. Therefore, chronic infections are usually treated with a maintenance therapy of multiple courses of administration to control the bacterial load in the airways^1^, which has been shown to increase the incidence of resistance to the applied antibiotics even further^19^.

While environmental *P. aeruginosa* strains can use a variety of resistance mechanisms^20^, virtually all aminoglycoside-resistant strains found in the CF lung show some degree of upregulation of the multi-drug resistance *mexXY* operon^21^ (**Fig. 1A**), which codes for an inner-membrane antiporter MexY and a periplasmic fusion protein MexX^22^ that mediate the export of a variety of different drugs, including most aminoglycosides^23^. Induction of the aminoglycoside response is primarily regulated by MexZ, a transcription repressor of *mexXY*^24^. In strains with a wild type *mexZ* allele, the activation of *mexXY* is caused by a decrease in translation efficiency, and not from a direct interaction with the drugs themselves, which allows resistance to be activated by a wide variety of stressors^25^. Loss of ribosomal function results in activation of anti-repressor ArmZ, due to loss of translational repression by a leader peptide^26^. ArmZ then binds and inactivates MexZ, eventually leading to the de-repression of *mexXY*. The outcome of drug treatments therefore depends on a lengthy process of MexXY activation that spans up to ~6 hours, and which only initiates when translation is already compromised^27^.

**Figure 1.**
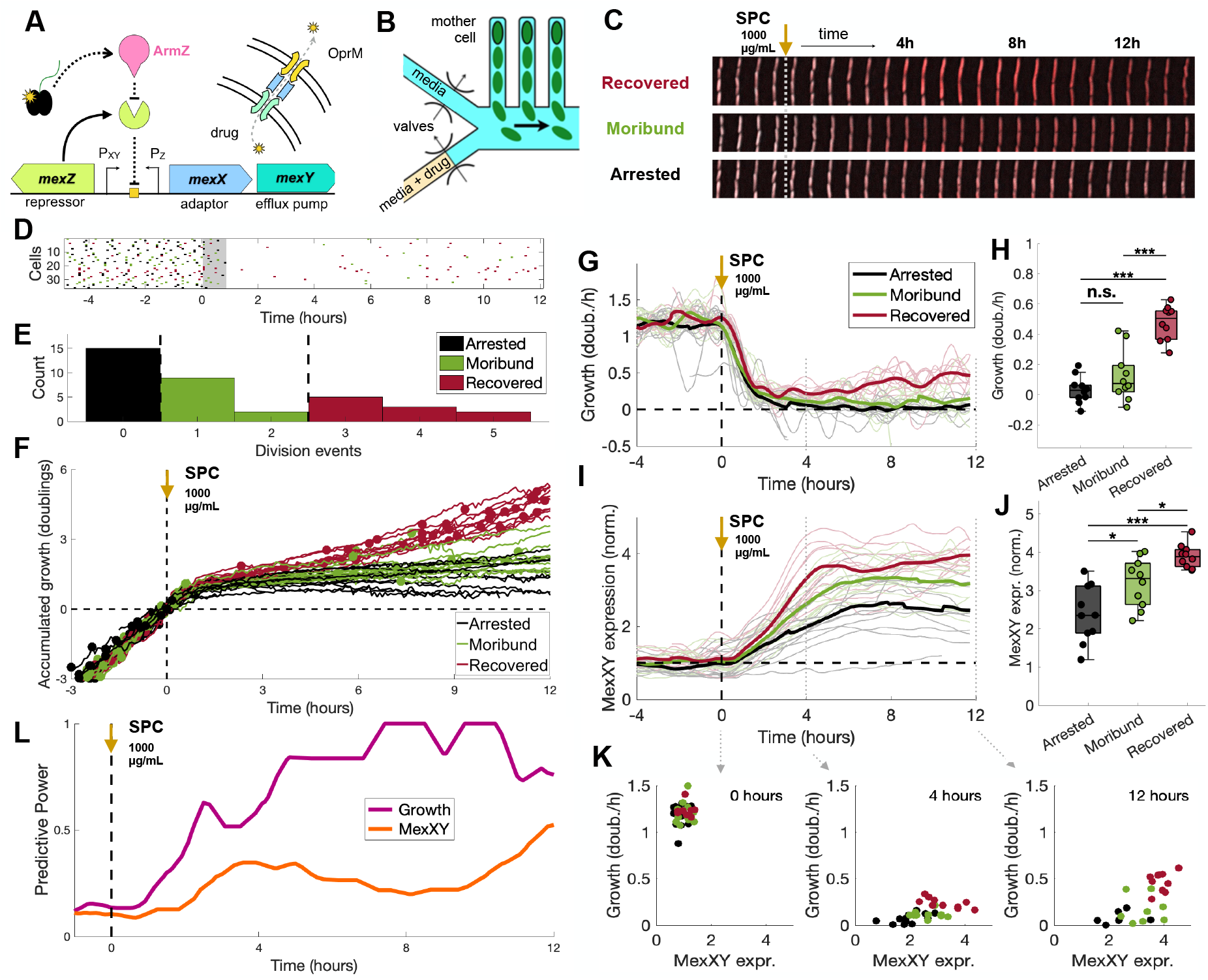
Abrupt exposure of single WT cells to spectinomycin shows heterogeneous response. **(A)** The MexXYZ resistance mechanism: aminoglycoside spectinomycin (SPC), a translation inhibitor, diffuses across the cell membrane and binds ribosome. The resulting decrease in translation efficiency induces expression of anti-repressor ArmZ, which binds repressor MexZ and releases expression of adaptor protein MexX and efflux pump MexY. The multidrug efflux system composed of MexX, MexY, and outer membrane protein OprM exports spectinomycin out of the cell. **(B)** Design of the microfluidic device, which places single cells in fixed locations as they go through division cycles. Valves allow on-chip rapid switching of media. WT cells carrying the native resistance mechanism with fluorescent reporters were exposed to a single step increase of 1000 μg/mL spectinomycin. **(C)** Time courses of individual cells representing the three different cell fates observed. Expression of *mexXY* is measured by a red fluorescence marker. **(D)** Cell division events for each mother cell throughout the experiment. The Y-axis represents each individual mother cell, and the dots denote division events colored by cell fate. Frequency of cell divisions decrease during the first ~90 min of drug exposure (highlighted). **(E)** Histogram showing number of division events in each cell after recovery from drug exposure. Cell fate was classified based on number of divisions: “arrested” if the cell did not divide, “moribund” if it divided 1 or 2 times, and “recovered” if it divided 3 or more times. **(F)** Accumulated growth in cell length for all cells observed, colored by cell fate and with division events indicated by dots. **(G)** Growth rate in each cell, with cell fate indicated by color. The thick line represents the average growth of cells with similar fate. **(H)** Final growth rates of each phenotype at the end of the experiment. The asterisks in each pairwise comparison denote a significance of *** p < 0.001, ** p < 0.01, and * p < 0.05 (**Methods**). **(I)** Expression of MexXY in each cell, with cell fate indicated by color. The thick line represents the average expression of cells with similar fate. **(J)** Final *mexXY* expression level of each phenotype at the end of the experiment. **(K)** MexXY levels and growth rate at three time points following exposure to spectinomycin, showing the distinct expression pattern of each cell fate. **(L)** Predictive power of cell growth and expression of MexY in determining recovery during early response. Cell growth predicts recovery within the first two hours following exposure, before predictions from MexXY. Pre-exposure growth and MexXY levels are not predictive of recovery.

Recent studies found that the single most common mutation observed in CF clinical samples is the loss of function of MexZ^28,29^, resulting in continuous expression of *mexXY*^30^. Despite being common in clinical samples, the evolution of *mexZ* mutants has not yet been reproduced *in vitro* in standard serial passage experiments^31,32^, and *mexZ* mutations do not appear to significantly increase the MIC of aminoglycosides^33^. However, these previous studies assessed resistance under steady drug concentrations, where MexXY is fully induced in wild type and *mexZ* mutants alike^30^. Therefore, we investigate the possibility that such regulatory mutations are beneficial during induction of the transcriptional response, before full expression is reached. We utilize single-cell and population-level assays to test the hypothesis that the continuous expression of *mexXY* in loss-of-function *mexZ* mutants bypasses the lengthy activation process of this resistance mechanism and allows a higher fraction of cells to survive rapid shifts in drug concentration. We propose a novel “Response Dynamics” AST assay, which quantifies the recovery of microbial populations following drug exposures and is complementary to classical MIC assays. We test this approach by profiling a collection of representative *P. aeruginosa* isolates, associating *mexZ* loss-of-function mutations with faster recoveries from exposures to aminoglycosides.

## Results

### Single-cell microfluidics shows heterogeneous response to aminoglycosides

To follow physiological changes in a population of single cells in real time as they respond to a step increase in drug concentrations, we developed a microfluidic device that combines mother-machine single-cell trapping with fast medium switching^2,34^ (**Fig. 1B, S1A**). A population of individual “mother cells” are each trapped at the closed end of short narrow chambers connected on their open side to a larger feeding channel where fresh minimal M63 medium supplemented with glucose, casamino acids, and thiamine is flowed. As the cells grow and divide, daughter cells are pushed into the feeding channel and are flushed away, keeping the mother cell in place through many generations. A system of valved inputs allows rapid switching between medium with and without drug.

To measure expression of the *mexXY* operon during responses to aminoglycosides, we have designed a promoter fusion placing a gene coding for mCherry under the regulation of the *mexXY* promoter, as well as fast folding GFP expressed from a constitutive P_*rpoD*_ promoter, in the same pMQ95 vector^35^ (**Methods, SI**). MexXY expression levels are calculated by normalizing the mCherry fluorescence signal by the GFP signal, which serves as a control and allows filtering of global effects on transcription. We have transformed this plasmid into the PA14 lab strain, together with motility Δ*flgK*/Δ*pilA* deletions needed to keep the bacteria in the chambers (referred here as WT). As an example of aminoglycoside antibiotic, we chose spectinomycin, which is known to induce strong MexXY responses^36^.

Following an abrupt and sustained exposure to 1000 μg/ml spectinomycin, which is close to the IC_50_ concentration where population-level growth is reduced to half, we observed wide ranges of growth rates and MexXY expression among single cells (**Fig. 1C**). All cells experienced an initial decrease in growth rate, and cell division ceased over the first 90 minutes following drug exposure (**Fig. 1D**). However, after a few hours many cells started to recover growth. Some of these growing cells sustained growth and resumed regular cell division, while others lingered in a state of slow growth and irregular division for the duration of the experiment. To compare these subpopulations, we categorized cell fates based on the number of division events after the first 90 minutes as Recovered (>3 divisions, n=10 cells), Moribund (1 or 2 divisions, n=11 cells), or Arrested (0 divisions, n=15 cells) (**Fig. 1E-H**). We next asked whether these different cell fates were dictated by the expression of MexXY at the onset of the drug. We followed the fluorescent reporters for MexXY in individual cells during the dynamic response and separated the cells according to their fate. While all cells showed some degree of MexXY upregulation during the response, recovered cells expressed MexXY the fastest, while moribund and arrested cells showed a slower response (**Fig. 1I-K, S1B, S2**). Recovered cells reached and sustained ~4-fold higher levels of MexXY expression, significantly above moribund or arrested cells. In general, induction of MexXY in recovered cells happened within 4 hours from drug exposure and preceded growth recovery, suggesting that MexXY is indeed necessary for cell survival^27^.

To understand whether early expression of resistance can predict recovery, we used an information gain approach to calculate the prediction power of gene expression or growth in the outcome of individual cells^2,37^. Recovery can be reasonably predicted from MexXY expression levels about four hours into the response, when recovered cells start to resume growth (**Fig. 1L**). However, recovery could not be predicted from initial levels of MexXY expression at the time of drug exposure. Therefore, small subpopulations can survive aggressive drug treatments even in homogeneous environments with no detectable pre-existing variations in growth or expression of resistance, and cell fate is determined only after drug exposure. Interestingly, cell growth itself is a better predictor of recovery, predicting cell fates even during the slow growth period following drug exposure. Therefore, it appears that cells with slightly higher levels of metabolic activity are poised to induce expression of MexXY and subsequently resume growth.

### Loss of MexZ repression increases cell survival upon drug exposure

We next investigated the role of MexZ regulation on single cell survival and the emergence of heterogeneity during the response to drug treatment. The effects of *mexZ* mutations on the fitness of *P. aeruginosa* during antibiotic treatments is not clear, as such mutations do not increase resistance by traditional MIC measures^33^. Although a deletion of the gene coding for repressor MexZ has been reported to increase MexXY expression in comparison to the wild type in the absence of the drug, fully induced MexXY levels in the presence of the drug are similar in the WT and *mexZ* mutant strains alike^30^. Therefore, differences in fitness are unlikely to arise during steady-state growth under drug exposure^31,32^, when resistance is fully induced. We then hypothesize that the continuous expression of *mexXY* in loss-of-function *mexZ* mutants bypasses the lengthy activation process of the resistance mechanism and allows a higher fraction of cells to survive rapid shifts in drug concentration. To test this hypothesis, we engineered a Δ*mexZ* strain with a clean deletion of the *mexZ* gene, derived from the same WT strain used in the microfluidic experiment above, as well as a Δ*mexY* strain with a deletion of the gene encoding the MexY efflux pump as a negative control. We then repeated the microfluidic experiment with these Δ*mexZ* and Δ*mexY* mutant strains.

In contrast to the heterogenous response observed in the wild-type strain, the Δ*mexZ* mutant maintained growth upon drug exposure, with all cells surviving (**Fig. 2A-C, S3**). Unlike WT cells, which experienced a decrease in growth following exposure prior to induction of MexXY, all Δ*mexZ* cells quickly equilibrated to a slightly lower steady-state growth rate in the presence of the drug. Meanwhile, unsurprisingly, all Δ*mexY* mutant cells quickly stopped growing following exposure, with no cells recovering growth. Thus, cell growth recovery depends on *mexXY* expression and the *mexZ* deletion indeed increase single-cell survival to drug exposures. Overall, phenotypic heterogeneity following drug exposure was exclusive to WT cells, and the fate of WT cells spanned the space between the phenotypes of the *mexXY* and *mexZ* mutants. Therefore, the emergence of heterogeneity depends on MexZ regulation of drug resistance and correlates with the noisy induction of resistance genes.

**Figure 2.**
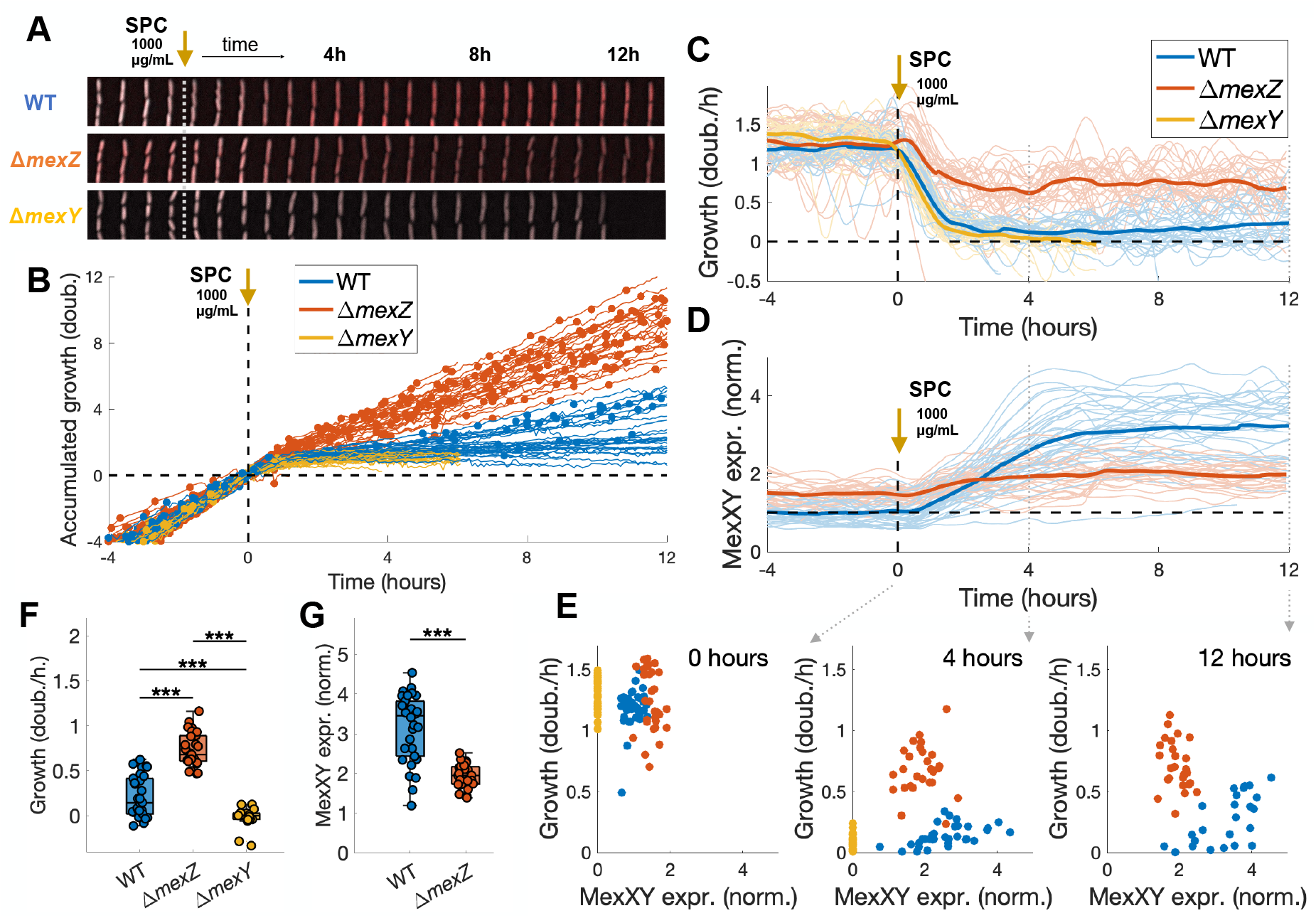
*mexZ* deletion increases single-cell survival to spectinomycin exposures. **(A)** Time courses of individual WT, Δ*mexZ*, and Δ*mexY* cells following an exposure to 1000 μg/mL spectinomycin in the microfluidic device. Expression of *mexXY* is measured by a red fluorescence marker. The Δ*mexZ* mutant has higher pre-drug *mexXY* expression levels and maintains cell growth and division following exposure. **(B)** Accumulated growth in cell length for WT, Δ*mexZ*, and Δ*mexY* cells, colored by genotype, with division events indicated by dots. **(C)** Growth rate in each cell, with genotype indicated by color. The thick line represents the average growth of cells within each genotype. **(D)** Expression of MexXY in each cell, with genotype indicated by color. The thick line represents the average expression of cells within each genotype. **(E)** MexXY levels and growth rate at three time points following exposure to spectinomycin, showing the distinct expression pattern of each genotype. **(F)** Final growth rates of each genotype at the end of the experiment. **(G)** Final *mexXY* expression level of WT and Δ*mexZ* cells at the end of the experiment. The asterisks in each pairwise comparison denote a significance of *** p < 0.001 (**Methods**).

Next, we compared the expression of MexXY between WT and Δ*mexZ* mutants (**Fig. 2D-G**). Δ*mexZ* mutants started with basal levels of MexXY ~2-fold higher than the WT in the absence of drug. After drug exposure, the slight decrease of growth rate in Δ*mexZ* cells corresponded to a slight increase of MexXY levels. Surprisingly, MexXY levels in Δ*mexZ* cells remained significantly below that reached by recovered WT cells (**Fig. S1C**). While Δ*mexZ* cells expressed sufficient levels of MexXY prior to the exposure and remained in a similar growth state in the presence of the drug, WT cells only induced MexXY expression after growth was decreased. Under slow growth conditions, fully induced MexXY in WT cells reaches higher levels than the Δ*mexZ* mutant^38^. This result is consistent with other studies where slow growth resulted in higher expression of resistance genes^39,40^. Taken together, these results suggest that sudden drug exposures take a significant toll on WT populations, with only a subpopulation of cells managing to induce MexXY-mediated resistance and recover growth (**Fig. S1D**). Therefore, the Δ*mexZ* mutant should have a fitness advantage over the WT in dynamic antibiotic exposure environments. Next, we test this hypothesis in competition assays.

### Loss of MexZ function confers a temporary advantage following drug exposure

Although Δ*mexZ* mutant cells have a higher survival rate than the WT during an abrupt exposure to spectinomycin, recovered WT cells that successfully induce MexXY can grow just as fast under sustained drug conditions. Following a sudden drug exposure, faster growing recovered cells tend to dominate the WT population over time, eventually bringing the WT population to a similar growth rate as the Δ*mexZ* mutant. Therefore, we predict that *mexZ* mutations bring only a temporary fitness advantage during sustained drug exposures, which should disappear as the WT population recovers growth. To test this prediction, we conducted competition experiments using a continuous culture device (chemostat), which allows the propagation of liquid cultures in carefully controlled environments for many generations (**Fig. 3A, S4, S5**). We built a device capable of measuring optical density (OD) and fluorescence, which we use to measure the size of the subpopulation of cells corresponding to the Δ*mexZ* strain, in which we introduced a constitutively-expressed mKO fluorescent marker^41^ (**Methods**). Therefore, our competition assays with mixed cultures of Δ*mexZ* and WT strains can determine the relative abundance of Δ*mexZ* cells in real time. In each experiment, we also verified the abundance of Δ*mexZ* and WT cells by counting CFUs at select time points.

**Figure 3.**
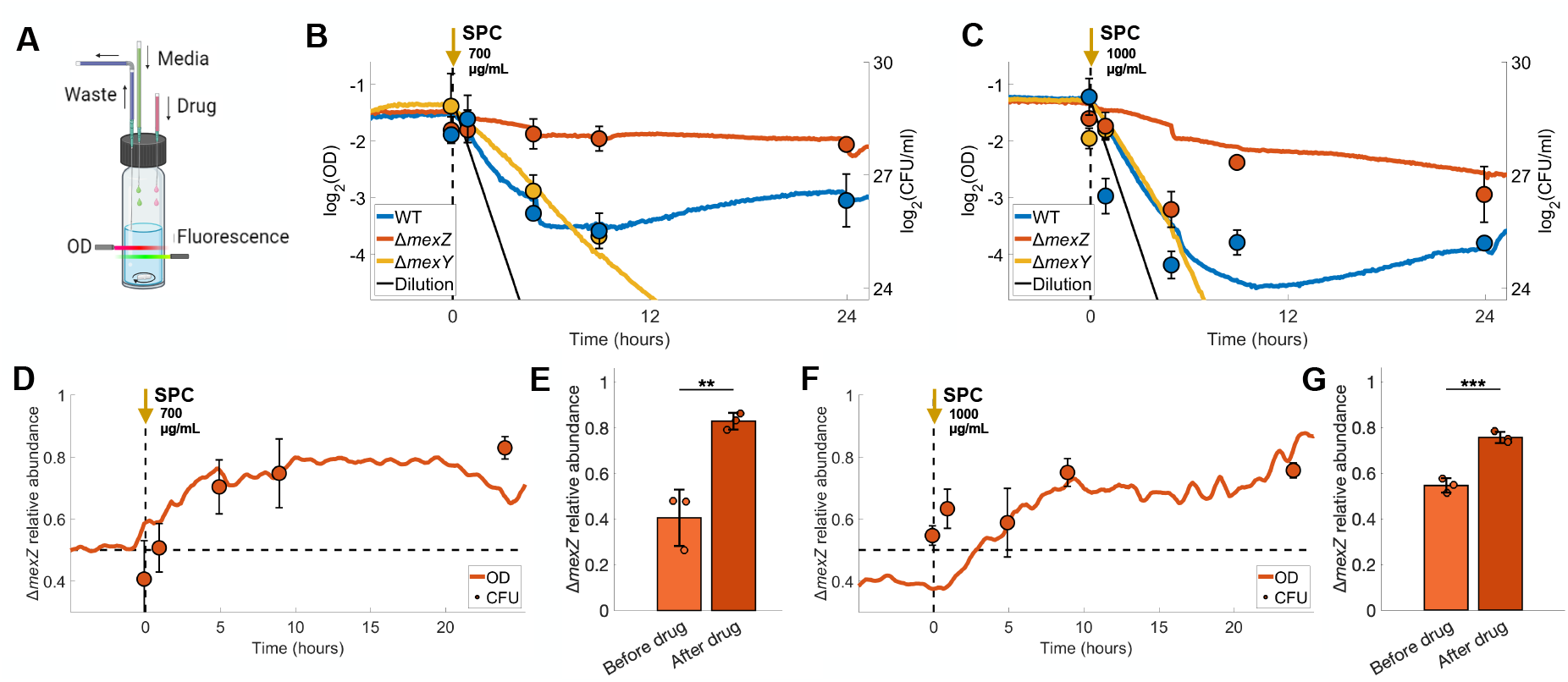
*mexZ* mutants have a temporary fitness advantage over the WT following drug exposure. **(A)** Continuous culture system (chemostat) dilutes culture with fresh media at a fixed rate, while keeping the culture at the same volume by removing spent media. OD and fluorescence are measured every 5 minutes. After growth is stabilized in fresh medium, drug is quickly added to the culture to the desired concentration, and medium is switched from fresh to drug medium for the duration of the experiment. **(B-C)** Monocultures of WT, Δ*mexZ*, and Δ*mexY* strains, exposed to **(B)** 700 and **(C)** 1000 μg/mL spectinomycin. Lines represent OD measurements in the chemostat. Dots represent the CFU counts of 3 plate replicates. **(D-G)** Δ*mexZ* and WT competition in mixed culture during **(D)** 700 and **(F)** 1000 μg/mL spectinomycin exposures. Lines represent the relative abundance of Δ*mexZ*, calculated from fluorescence and OD measurements in the chemostat (**Methods**). Dots represent the relative abundance of Δ*mexZ* in CFU counts of 3 plate replicates. Bar plots indicate the relative abundance of the Δ*mexZ* strain at the moment of exposure to **(E)** 700 and **(G)** 1000 μg/mL spectinomycin and at the end of the experiment. The asterisks in each pairwise comparison denote a significance of *** p < 0.001 and ** p < 0.01 (**Methods**).

To determine how growing populations respond to sudden exposure to antibiotics, we started by growing WT, Δ*mexZ*, and Δ*mexY* as monocultures and exposed them to a single, sustained step increase in drug concentration (**Fig. 3BC**). We ran these experiments at a relatively low dilution rate of 0.6 hour^-1^, which allowed each strain to stabilize growth at an OD of ~0.4 in the absence of drug, still below stationary phase. Following an exposure to 1000 μg/ml spectinomycin, both WT and Δ*mexY* populations were initially quickly reduced a rate close to the dilution rate, which would be expected if all cells were arrested. However, while Δ*mexY* was subsequently washed out of the culture, the WT started to recover growth after ~5 hours. Meanwhile, the Δ*mexZ* culture did not experience a sharp decrease in OD and slowly adjusted to growth at a slightly lower OD in the presence of the drug. Both WT and Δ*mexZ* were slowly converging to a similar OD towards the end of the experiment at 24 hrs (the experiment was stopped at 24 hours to prevent the emergence of mutants). We saw a similar pattern with an exposure to 700 μg/ml spectinomycin, where the WT was initially diluted but recovered after ~5 hours, while the Δ*mexZ* mutant did not experience drastic changes in OD. CFU counts largely followed OD measurements, despite slight differences due to inviable cells impacting OD but not CFU measures. Overall, these experiments show that loss of *mexZ* can shorten the population-level recovery to abrupt drug exposures.

Next, we competed the Δ*mexZ* mutant with the WT in mixed cultures to determine changes in their relative abundance during drug exposures (**Fig. 3D-G**). In the absence of drug, both strains were stable at ~50% abundance, indicating that *mexZ* mutations do not bring significant fitness costs under our experimental conditions. After exposures to 700 and 1000 μg/ml spectinomycin, the relative abundance of Δ*mexZ* mutants in the population grew to ~80% over 5 to 10 hours, but then stabilized at these values. These relative abundances were also confirmed by CFU counts. Therefore, *mexZ* mutations indeed provide a temporary fitness advantage in direct competition with the WT in the presence of spectinomycin. These results suggest that such regulatory mutations are expected to rise in *P. aeruginosa* populations frequently subjected to antibiotic treatments.

### “Response Dynamics” assay to measure population recovery from drug exposures

Since *mexZ* mutations do not provide a clear fitness advantage during steady-state growth in the presence of the antibiotic, the advantage such mutations provide during sudden drug exposures is not captured by traditional MIC measures of resistance obtained by growing microbial cultures under steady drug conditions. Therefore, we developed a “Response Dynamics” assay to profile antibiotic resistance in a dynamic context^42^. In this assay, growing liquid cultures are exposed to a sharp and sustained increase in drug concentration, which allows us to quantify population-level recovery time (**Fig. 4A, S6**). The arrest of a large fraction of a growing bacterial culture following a drug exposure is manifested as a pause (or decrease) in population-level growth, during the time it takes for the subpopulation of recovered cells to grow past the background of arrested cells. This pause in growth accounts for the contributions of both cell death and slow growth. We can then measure the new growth rate upon the population re-initiating growth.

**Figure 4.**
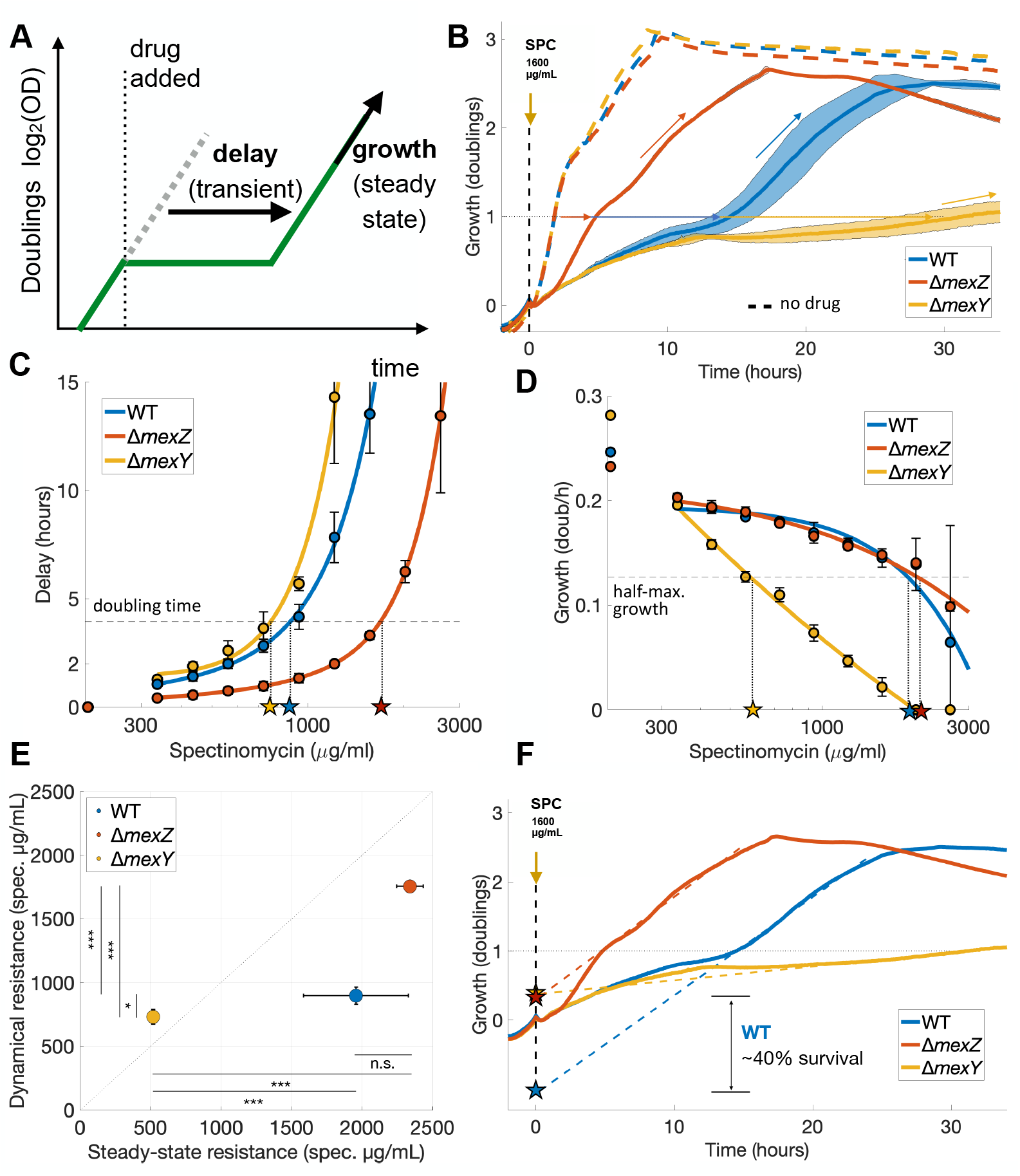
Response Dynamics assay: steady-state and dynamic resistances describe the antibiotic resistance profile of *P. aeruginosa* in a dynamic context. **(A)** Sudden exposures to antibiotics cause a pause in population growth, as a significant fraction of the population is arrested. Growth resumes once a “recovered” subpopulation of cells that successfully induced the drug response grows past the background of arrested cells. The Response Dynamics assay uses this pause in growth as a measure of population recovery from drug exposures. **(B)** Growth of WT, Δ*mexZ*, and Δ*mexY* strains upon exposure to a step increase of 1600 μg/mL spectinomycin (at time zero). Horizontal arrows indicate the delay in the time taken to reach one doubling in comparison to growth in the absence of drug. Inclined arrows indicate the steady-state growth reached after population recovery. **(C)** Delays in recovery for each strain as a function of the spectinomycin dose used in the exposure. The horizontal line indicates the threshold of one doubling time, measured during exponential growth in the absence of drug. We calculate *Dynamic Resistance* as the drug concentration that causes a delay in recovery equal to this threshold, indicated by stars. Dynamic resistance quantifies the capacity for quick recovery following drug exposure. **(D)** Steady-state growth of each strain as a function of the spectinomycin dose used in the exposure. The horizontal line indicates the threshold of half of the maximum growth rate, measured during exponential growth in the absence of drug. We calculate *Steady-state Resistance* as the drug concentration that reduces the steady-state growth rate to this threshold, indicated by stars. Steady-state resistance measures the capacity for growth in the presence of drug, similar to the IC_50_ and other traditional measures of resistance such as MIC. **(E)** Dynamic and Steady-state resistances for the WT, Δ*mexZ*, and Δ*mexY* strains. *mexZ* deletion increases Dynamic resistance without affecting Steady-state resistance. The asterisks in each pairwise comparison denote a significance of *** p < 0.001, ** p < 0.01, and * p < 0.05 (**Methods**). **(F)** For the heterogeneous drug response of the WT strain, an “effective survival rate” can be calculated by projecting steady-state growth of the recovered population to the time of drug exposure. The responses of the Δ*mexZ* and Δ*mexY* strains, which are homogeneous albeit at different growth rates, project steady-state growth back to a point close to the population at the time of drug exposure.

In this assay, we expose an array of mid-log phase bacterial cultures to shifts in a large span of drug concentrations, and measure both the duration of the pause in growth following exposure and the steady-state growth rate after population growth is resumed. We extract from this assay two complementary measures of resistance: 1) *Dynamic Resistance*, which is the drug concentration that introduces a delay equivalent to the pre-drug doubling time in population-level recovery time from drug exposure, and 2) *Steady-State Resistance*, which is the drug concentration that halves the growth rate of the recovered population (similar to the IC_50_ concentration, which correlates with MIC). We express these measures of resistance in units of drug concentration, in line with other typical measures. For instance, high dynamic resistance means that a large drug dose is necessary to significantly impact the growth of single cells following exposure, and therefore this population can recover quickly from exposures to lower doses. The thresholds used to calculate these measures of resistance – the pre-drug doubling time and growth rate – were chosen to account for differences in cell growth when comparing different strains. Thus, our assay captures the dynamic aspect of antibiotic responses, which is critical for resistance and is not measured by MIC assays.

To verify that the Response Dynamics assay can capture the fitness advantage of *mexZ* mutations, we performed the assay in the WT, Δ*mexZ*, and Δ*mexY* strains. Following exposures to high drug concentrations close to the MIC, we observed long delays in growth recovery for WT cultures, while Δ*mexZ* cultures kept growing without interruptions and Δ*mexY* cultures failed to recover growth (**Fig. 4B, S7**). However, the steady-state growth rate following recovery was still similar between WT and Δ*mexZ*. Therefore, the heterogeneous behavior of WT cells following drug exposures translates into reliable population-level behaviors that can be quantified.

Next, we calculated steady-state and dynamic resistances. For all strains, both the speed of recovery from drug exposures (**Fig. 4C**) and the steady-state growth rate (**Fig. 4D**) decline as a function of the spectinomycin dose. While steady-state growth declines rapidly with drug dose for the Δ*mexY* strain, WT and Δ*mexZ* can sustain growth under much higher doses, with the growth rate declining in a very similar manner for both strains. Therefore, the presence of a functional MexXY affords resistance regardless of regulation, giving WT and Δ*mexZ* the same steady-state resistance, which is consistent with the similar MICs reported in the literature^33^ (**Fig. 4E**). On the other hand, while the Δ*mexZ* strain only showed delays in recovery at high drug doses, the WT showed significant delays at much lower drug doses, only slightly higher than the Δ*mexY* strain. Therefore, the resulting lower dynamic resistance of the WT in comparison to Δ*mexZ* captures the WT’s faltering population growth caused by its heterogeneous drug response. In fact, an “effective survival” rate can be calculated by extrapolating the steady-state growth of the recovered population back to the time of drug exposure, showing a ~40% survival rate for the Δ*mexZ* mutant that is consistent with the heterogeneity observed in single-cell microfluidic experiments (**Fig. 4F**). However, since the WT strain shows a broad distribution of growth rates following drug exposure, and steady-state growth is dominated by the fastest-growing cells, this is an imprecise measure of survival that is less useful in calculating dynamic resistance.

Although steady-state and dynamic resistances measure different properties relating to antibiotic resistance, they are not completely independent, as they depend on the action of the same resistance mechanism. Steady-state resistance relates to the efficiency of resistance mechanisms, measuring their ability to support growth at higher drug concentrations. On the other hand, dynamic resistance relates to the regulation of resistance mechanisms, and measures the ability of cells to quickly activate the appropriate responses and recover growth upon sudden drug exposures. Therefore, a more efficient resistance mechanism, capable of clearing drug at a higher rate, will increase both measures of resistance. However, slower responses will result in lower dynamic resistance, without necessarily changing steady-state resistance. This is evident by the decrease in dynamic resistance caused by the presence of MexZ regulation, which does not change steady-state resistance (**Fig. 4E**). Meanwhile, steady-state and dynamic resistances are similar for Δ*mexZ*, where resistance is constitutively expressed, since growth and speed of recovery start decreasing at similar drug doses. Steady-state and dynamic resistances are also similar for Δ*mexY*, where resistance is absent, as its survival to drug exposures does not depend on the induction of the MexXY response.

### *mexZ* truncations are associated with high dynamic resistance in a diverse collection of isolates

Although our previous experiments show that a *mexZ* deletion accelerates recovery from spectinomycin exposures in a PA14 lab strain, it was still unclear how commonly found *mexZ* mutations affect antibiotic resistance in natural strains. To answer this question, we assembled a representative panel of 49 *P. aeruginosa* environmental and clinical specimens reflecting the organism’s diversity, obtained from the International Pseudomonas Consortium Database (IPCD)^12,43–45^. This set includes reference genomes, transmissible strains, sequential CF isolates, strains with specific virulence characteristics, and strains that represent serotypes, genotypes, or geographic diversity (**Fig. 5A, S8, Table 1**). Importantly, this panel includes 12 strains with truncations or large indels in the *mexZ* gene, which is otherwise well conserved (**Fig. 5B**). We then used our Response Dynamics assay to measure steady-state and dynamic resistances to spectinomycin in this collection of representative strains, finding a wide diversity of resistance profiles to this drug (**Fig. 5C, S9-13**). We find that in most strains dynamic resistance is lower than steady-state resistance, suggesting heterogeneous responses with delayed recovery from drug exposures, as was seen with our WT strain. However, strains with *mexZ* truncations tend to be closer to the diagonal where dynamic resistance is similar to steady-state resistance, which is consistent with constitutive *mexXY* expression and fast recoveries, as seen with our Δ*mexZ* mutant. Next, we quantitatively assess these trends.

**Table 1.** Panel of *P. aeruginosa* specimens reflecting the organism’s diversity. Strains were imported from the IPCD collection, which includes clinical and environmental samples. We noted any *mexZ* mutations in relation to the gene’s consensus sequence. *mexZ* variants were classified as “truncations” in case of large indels or frameshift mutations that resulted in the loss of a significant part of the gene. Alignment scores of *mexZ* variants with the consensus sequence are also included. Dynamic and steady-state resistances for each strain were measured by the Response Dynamics assay (**Methods**). Shaded cells indicate strains that did not grow in our culture medium formulation and were therefore excluded from the analysis.

| IPCD | Name | Environ. | MexZ mutations | Trunc. | Score | Dynamic resistance<br>($\mu\text{g/ml Spec.}$) | Steady-state resistance<br>($\mu\text{g/ml Spec.}$) |
| --- | --- | --- | --- | --- | --- | --- | --- |
| 80 | AMT0060-1 | CF | - | 0 | 466.00 | 1068.00 | 1491.52 |
| 81 | AMT0060-2 | CF | - | 0 | 466.00 | 1231.11 | 743.30 |
| 82 | AMT00060-3 | CF | - | 0 | 466.00 | 1470.47 | 732.81 |
| 83 | PAK | Human | - | 0 | 466.00 | 1083.28 | 1953.94 |
| 84 | IST27 | CF | - | 0 | 466.00 | 1023.41 | 1872.38 |
| 85 | IST27N | CF | - | 0 | 466.00 | 654.05 | 913.42 |
| 86 | 968333S | Human | I141N | 0 | 463.33 |  |  |
| 87 | 679 | Human | - | 0 | 466.00 | 194.04 | 333.00 |
| 88 | NH57388A | CF | truncation R153 (frameshift insertion) | 1 | 326.66 |  |  |
| 89 | 1709-12 | Human | - | 0 | 466.00 | 2500.00 | 2500.00 |
| 91 | Jpn1563 | Lake | - | 0 | 466.00 | 539.85 | 1016.17 |
| 92 | LMG14084 | Water | - | 0 | 466.00 | 70.73 | 143.97 |
| 93 | Pr335 | Hospital | L128M | 0 | 465.33 | 365.20 | 622.31 |
| 94 | U018A | CF | - | 0 | 466.00 | 613.52 | 1106.63 |
| 95 | CPHL9433 | Plant | L138R \| T177S | 0 | 462.33 | 143.97 | 217.40 |
| 252 | AUS226 | River | L138R \| E146A \| F192L \| L196I | 0 | 457.67 | 307.93 | 786.78 |
| 253 | AUS227 | River | L138R | 0 | 463.33 | 162.43 | 468.33 |
| 258 | AUS232 | Animal | L138R \| L196I | 0 | 462.33 | 325.94 | 775.68 |
| 300 | AUS327 | CF | deletion I161_L180 | 1 | 368.66 |  |  |
| 349 | AUS089 | CF | truncation N76 (frameshift deletion) | 1 | 146.00 | 252.04 | 194.75 |
| 436 | AUS485 | Animal | P80Q \| I84L \| L138R \| N186S \| L196I | 0 | 455.67 | 312.36 | 485.30 |
| 448 | AUS511 | River | L138R | 0 | 463.33 | 276.79 | 587.91 |
| 450 | AUS526 | Home | - | 0 | 466.00 | 838.75 | 2500.00 |
| 543 | AMT0005-138 | CF | truncation K4 (frameshift deletion) | 1 | -34.33 | 1523.66 | 345.01 |
| 573 | AMT0071-76 | CF | V29E \| L37Q \| L138R \| N186S | 0 | 456.33 | 415.06 | 211.31 |
| 1103 | AUS23 | CF | A38T | 0 | 464.33 | 1098.79 | 1138.54 |
| 1256 | C3719 | CF | truncation 172 (frameshift deletion) | 1 | 365.00 | 781.21 | 1379.37 |
| 1257 | AES-1R | CF | Y52_K53 ins KN | 0 | 460.67 | 2500.00 | 2039.06 |
| 1258 | AUS52 | CF | truncation E114 (frameshift insertion) | 1 | 228.66 |  |  |
| 1259 | AA2 | CF | I141N | 0 | 463.33 | 345.01 | 980.69 |
| 1260 | AMT0023-30 | CF | - | 0 | 466.00 | 826.91 | 1556.49 |
| 1261 | AMT0023-34 | CF | truncation A61 (frameshift deletion) | 1 | 105.66 | 1819.90 | 1512.87 |
| 1262 | CHA | CF | L25P \| L138R | 0 | 460.33 | 697.25 | 1502.16 |
| 1264 | 39016 | Human | - | 0 | 466.00 | 2500.00 | 2268.43 |
| 1266 | Mi162-2 | Human | R87P \| G89S | 0 | 460.00 | 933.10 | 1293.90 |
| 1268 | 15108-1 | Human | - | 0 | 466.00 | 485.27 | 732.81 |
| 1269 | 13121-1 | Human | truncation 146 (frameshift insertion) | 1 | 308.67 | 2500.00 | 2500.00 |
| 1270 | 39177 | Human | L128M | 0 | 465.33 | 349.95 | 809.47 |
| 1271 | KK1 | CF | - | 0 | 466.00 | 532.27 | 543.74 |
| **1272** | A5803 | Human | - | 0 | 466.00 | 378.43 | 521.04 |
| **1273** | TBCF10839 | CF | - | 0 | 466.00 | 248.81 | 697.25 |
| **1274** | 57P31PA | Human | - | 0 | 466.00 | 869.09 | 1926.37 |
| **1360** | AL430 | CF | truncation R87 (large indel) | 1 | 174.33 |  |  |
| **1370** | AL199 | CF | truncation S198 (large indel) | 1 | 436.00 | 1832.88 | 2500.00 |
| **1423** | BJ4 | CF | V156I \| truncation 195 (large indel) | 1 | 427.00 | 1075.61 | 707.23 |
| **1432** | SC1 | CF | L138R \| N186S \| L196I | 0 | 460.33 | 1091.01 | 1122.47 |
| **1439** | Zw73-1 | CF | truncation 54 (frameshift insertion) | 1 | 95.33 | 1419.14 | 1601.38 |
|  | PAO1 | Lab strain | - | 0 | 466.00 | 571.44 | 1389.20 |
|  | PA14 | Lab strain | L138R \| N186S | 0 | 461.33 | 738.04 | 1819.90 |

**Figure 5.**
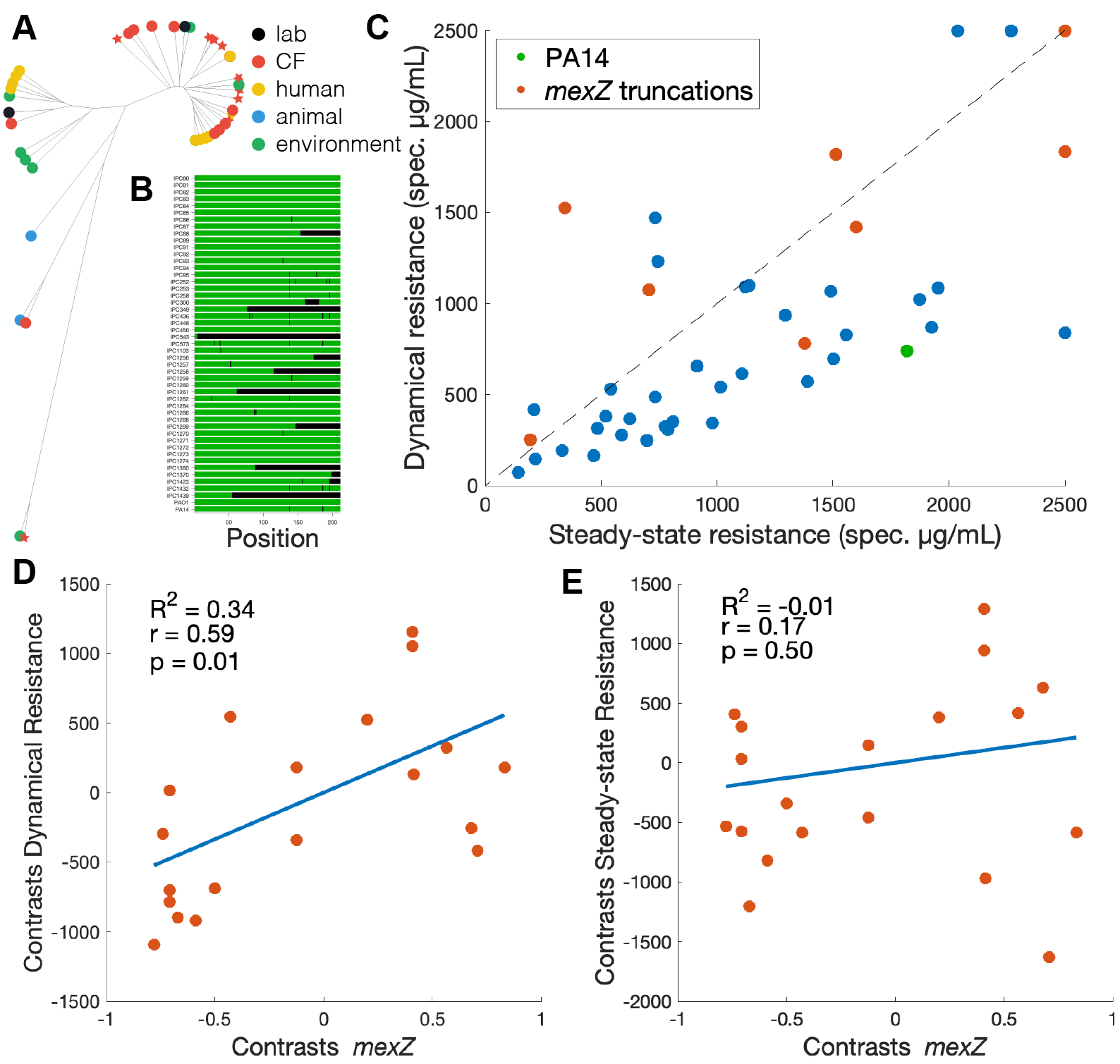
*mexZ* truncations are associated with high dynamic resistance in a panel of representative *P. aeruginosa* isolates. **(A)** Phylogenetic tree of *P. aeruginosa* specimens used in this study. This panel includes samples from diverse environments and represents the diversity of the organism. Strains with *mexZ* truncations are denoted by stars. **(B)** Mutations in the *mexZ* gene for each of the strains, in comparison with the consensus sequence. Other than the presence of truncations or large indels, the gene is well conserved. **(C)** Dynamic and steady-state resistances were measured for each of the strains with the Response Dynamics assay. Most strains with functional variants of MexZ show dynamic resistance lower than steady-state resistance, suggesting a heterogeneous response. Strains with *mexZ* truncations are closer to the diagonal where dynamic and steady-state resistances are similar. **(D)** Linear regression shows strong and significant correlation between the presence of *mexZ* truncations and dynamic resistance. We used phylogenetic independent contrasts to account for the underlying phylogenetic structure of the *P. aeruginosa* collection. **(E)** Linear regression shows no correlation between *mexZ* mutations and steady-state resistance.

To associate antibiotic resistance measured in our Response Dynamics assay with *mexZ* variants found in our collection of strains, we perform trait comparative analyses^46,47^ using phylogenetic independent contrasts (PIC)^48^, which account for the underlying phylogenetic structure of the *P. aeruginosa* collection. We obtained the phylogenetic tree of our collection of strains using 7 housekeeping genes commonly used for multi-locus sequence typing (MLST) (**Methods**). We find that *mexZ* truncations are interspersed throughout the tree (**Fig. 5A**), indicating that these mutations are frequently present in genetically distant strains, possibly conferring similar phenotypes. We then used this tree to calculate contrasts of dynamic and steady-state resistances, as well as contrasts relating to the presence of *mexZ* truncations, which can then be used for linear regressions to establish correlations (**Methods**). Defining *mexZ* truncations as a binary trait (presence/absence) results in several contrasts equal to zero, which are then discarded from the analysis. As an alternative, to increase the number of non-zero contrasts, we also calculated contrasts using the phylogenetic distance of each *mexZ* variant from the consensus *mexZ* sequence, which is a continuous trait. However, this did not result in better correlations (**Fig. S14**). Non-truncated variants of *mexZ* typically differ by only a few SNPs that do not necessarily contribute to its function^49^, and thus considering such differences added little information to the analysis. Therefore, we proceeded by classifying *mexZ* variants by the presence or absence of truncations or large indels.

We then used these contrasts to correlate the presence of *mexZ* truncations to the spectinomycin resistance phenotypes measured with our Response Dynamics assay in our strain collection. We found strong and significant correlation between the presence of *mexZ* truncations and dynamic resistance (*r* = 0.59, *p* = 0.01), which was able to explain 34% of the variance in the data (**Fig. 5D**). Conversely, we found no significant correlation between *mexZ* truncations and steady-state resistance (*r* = 0.17, *p* = 0.50) (**Fig. 5E**). These results are consistent with our experiments with the Δ*mexZ* lab strain, and show that that *mexZ* truncations disrupting MexZ function accelerate recovery from exposures to spectinomycin also in clinical strains. Therefore, our Response Dynamics assay quantifies an important aspect of antibiotic resistance that is missing from traditional measures, and can be used for a more complete profiling of clinical strains.

## Discussion

We have shown that the most common mutation found in clinical *P. aeruginosa* samples from individuals with chronic infections and continual exposure to antibiotics, a loss of function of repressor MexZ^28,29^, improves antibiotic resistance by increasing single-cell survival and shortening population recovery from abrupt exposures to the drug. This fitness advantage of *mexZ* mutations relates to the dynamics and heterogeneity of drug responses, which are frequently overlooked and are not captured by most standard methods of antibiotic susceptibility testing (AST). This concept does not fall neatly into the currently used terminology describing the many ways microbes can evade antibiotic treatments. The coexistence of growing and arrested bacterial cells in the presence of antibiotics is usually attributed to heteroresistance – as opposed to persistence, where subpopulations survive under the antibiotic without growing^50^. The underlying mechanisms of heteroresistance are often attributed to some pre-existing heterogeneity of resistance phenotypes in the population (e.g., stochastic genetic switches), which is then selected upon by the drug^8,51^. Here, we report initially homogeneous microbial populations that develop phenotypic diversity only after antibiotic exposures, a conclusion we draw because recovery depends on the expression of inducible mechanisms. This heterogeneity results from induction of resistance being hindered by the effects of the drug itself, which leads to either cell survival or death/arrest, depending on whether sufficient expression of resistance genes is quickly achieved.

The emergence of heterogeneity during the induction of antibiotic responses is a general phenomenon and has been observed for different drug classes and model systems^2,52^ (**Fig. S6**). However, single-cell and population-level behaviors can differ according to the specificities of each mechanism of drug action and antibiotic resistance^53^. At the population level, decreases in OD are frequently observed when the antibiotic causes cell lysis. At the single cell level, heterogeneity can present as either well-defined cell fates or a more continuous distribution of phenotypes. Here, the *P. aeruginosa* aminoglycoside response resulted in a wide distribution of growth rates and MexXY expression levels in WT cells, spanning the space between the uniform responses in the Δ*mexY* strain, where there is no resistance, and the Δ*mexZ* strain, where resistance is constitutively expressed. Contrastingly, the heterogeneity in the *E. coli* tetracycline response has been reported to quickly resolve into either cell arrest or full growth recovery^2^. Unlike the MexXY resistance mechanism, which has low drug specificity^22^ and is slowly induced^27^ within a low dynamic range^25^, the *tet* resistance mechanism is quickly induced to high levels upon drug exposure and provides resistance to drug concentrations ~100-fold higher in comparison to sensitive cells^2^. Therefore, it is likely that fluctuations in *tet* expression have a more dramatic effect on the survival of *E. coli* cells exposed to tetracycline. Nevertheless, regardless of such differences, heterogeneity and the resulting population effects are likely to be general to any scenario where cells need to quickly transition to an alternative phenotype to sustain growth in the presence of a stressor^52^.

To assess the resistance profile of bacterial samples, resistance is typically defined by estimating the minimum inhibitory concentration (MIC) required to inhibit the growth of bulk populations in the presence of antibiotics. However, in recent years, it has become clear that populations that are classified as susceptible by their MIC value may still harbor subpopulations that survive antibiotic treatment^3,51,54^, resulting in discrepancies between resistance profiling in the lab and the outcomes of clinical treatments^55^. While clinical antibiotic treatments are generally well designed to completely clear most infections, short courses of antibiotics such as intravenous or inhalation can be delivered within minutes and result in sharp increases of drug bioavailability at the site of infections^16,56^. Therefore, our Response Dynamics AST assay will be relevant in detecting strains with improved recovery to drug exposures in situations where drug delivery happens at shorter timescales than the induction of antibiotic responses. We note that we use the strain’s growth rate in the absence of drug to define both the threshold in steady-state growth used to calculate steady-state resistance (half of the pre-drug growth rate) and the threshold in delay of population recovery used to calculate dynamic resistance (pre-drug doubling time). Therefore, our assay can be used to analyze strains with different growth rates and can be performed in different media conditions. As we have shown with our analysis of a collection of *P. aeruginosa* strains, this assay captures gains in resistance that overlooked by traditional AST methods, and therefore has the potential to identify a new class of regulatory mutations that affect antibiotic resistance are common in clinical settings.

These results help explain how regulatory mutations play a critical role in the evolution of microbes in rapidly changing environments, conferring evolvability to cell responses and allowing quick adaptation across different ecological niches. Regulatory mutations, particularly disruptions of transcriptional repression, are much more easily achieved by evolution than the specific point mutations necessary to refine resistance proteins^57,58^. In particular, constitutive expression of broad spectrum efflux mechanisms like MexXY-OprM can confer dynamic resistance against a variety of stresses, such as multiple classes of antibiotics and ROS, which are released in neutrophil attacks^59,60^. Therefore, we expect that regulatory mutations increasing microbial survival to drug exposures are widespread in clinical settings. By maintaining population size and growth during antibiotic treatments, microbes can better evade clearance by the immune system^61^, increase tissue invasiveness^62^, and are more likely to acquire further mutations increasing resistance^63–66^.

## Materials and methods

### Media, drugs, and strains

All experiments were conducted in a modified M63 minimal medium (2 g/L (NH_4_)_2_SO_4_, 13.6 g/L KH_2_PO_4_, 0.5 mg/L FeSO_4_·7H_2_O) supplemented with 2 g/L glucose, 1 g/L casamino acids, 0.12 mg/L MgSO4, and 0.5 mg/L thiamine. Spectinomycin solutions were freshly made from powder stocks (Gold Bio). Spectinomycin was chosen for these experiments because it is reported to cause high induction of MexXY^36^. All strains used in this study were derived from *P. aeruginosa* PA14. The PA14 Δ*flgK*Δ*pilA* strain used in the microfluidic assays was a gift from the O’Toole lab. The deletion of flagellum hook-associated gene *flgK* and type-4 pilus associated gene *pilA* hinders cell motility to keep cells in place within the device. The PA14 strain carrying 2 chromosomal insertions of mKO, used in the continuous culture experiments, was a gift from the Nadell lab. A detailed description of our strains is provided in the Supplemental Information.

### Microfluidic experiments

*P. aeruginosa* cultures were grown overnight in LB, then inoculated into the device and forced into the cell channels by centrifugation. Immediately after inoculation, the microfluidic device was mounted on a Nikon TE2000-E microscope with Perfect Focus, a full incubation chamber, and a CoolSnapHQ Monochrome camera. Cells were allowed to equilibrate in the device for 12 hours before imaging. Image acquisition was performed at 37°C using Nikon NIS-Elements software. Exposures were done at very low illumination intensities to avoid light toxicity, with 2×2 binning for mCherry fluorescence but no binning for GFP fluorescence.

For each *P. aeruginosa* strain, we followed ~40 cells in the microfluidic device, recording cell sizes and fluorescence intensities every 5 min, over 20 hours. Cells were grown in fresh medium until growth was stabilized, then exposed to 1000 μg/mL of spectinomycin. We picked a drug concentration which impairs growth, close to the IC_50_ concentration where population-level growth is reduced to half. Gene expression was measured using a reporter plasmid expressing fluorescent reporters sfGFP (5.6 min maturation time) and mCherry (15 min maturation time) from the native promoters of RpoD and MexY, respectively. All data analysis was based on a custom Matlab image-processing pipeline. For each image, the top cell in each channel was identified and their length and mean fluorescence intensity was calculated. Cell division events were identified by looking for instances where a cell’s length dropped to less than 70% of its previous value (**Fig. S1A**). Growth rate was measured in doublings/hour by taking the derivative of log_2_(cell length).

### Significance in pairwise comparisons

Assumption of normal distributions in the data were checked using the Anderson-Darling test. For pairwise comparisons between normally distributed data, significance was determined using a two-way t-test with an alpha value of 0.05. Otherwise, we used a Mann-Whitney U-test. For multiple comparisons, ANOVA was performed using the Dunn–Šidák correction. The asterisks denote *** p < 0.001, ** p < 0.01, and * p < 0.05.

### Predictive power of an attribute

At the end of an experiment with *N* cells, the outcomes are divided into *N*_1_ recovered cells and *N*_2_ non-recovered cells (*N*_1_ + *N*_2_ = *N*), and the total information entropy of this system is

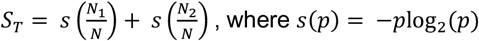

At time *t*, for given value *x*_*i*_ of an attribute *x* of the cell (MexXY expression, for instance), we divide the population into two subpopulations: *N*_*A*_ cells where *x* ≤ *x*_*i*_, and *N*_’_ cells where *x* > *x*_*i*_. Considering the outcome, the information entropy within each subpopulation at time *t* is

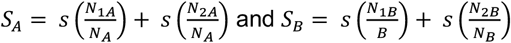

The information gain of splitting of the original population at *x*_*i*_ is then defined, as

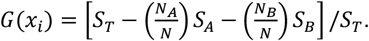

If each subpopulation at time *t* shows roughly the same distribution of outcomes as the final population, then *S*_*A*_ ≈ *S*_*B*_ ≈ *S*_T_ and *G* ≈ 0. However, if the population at time *t* is split along the *x*_*i*_ value that separate between recovered and non-recovered cells, then *S*_*A*_ ≈ *S*_*B*_ ≈ 0 and *G* ≈ 1. We define the predictive power of attribute *x* at time *t* as the gain obtained from the best split of the population, at the *x*_*i*_ value that maximizes *G*.

### Continuous culture experiments

Experiments were run in a custom-made device, which is further described in the Supplemental Information. Glass vials were coated with Sigmacote (Sigma) to prevent biofilm growth. Vials, vial heads, tubing, and all connections were autoclaved before assembly and after each experiment. Complete sterilization was ensured by running 10% bleach, 70% ethanol, and sterile water consecutively through all tubing and connections prior to each experiment. Experiments were carried out at 37 °C in sterile media. Vials were filled with 15 ml of culture media and used to blank OD measurements. Drug media consisted of fresh medium with spectinomycin added to either 700 or 1000 μg/ml, mixed until completely dissolved in solution, and filter sterilized into the container used for the experiment. To achieve an abrupt switch to drug in the chemostat, cultures were spiked with a drug stock to the desired concentration and the media was immediately swapped from fresh medium to medium with spectinomycin. Cultures were constantly diluted at 0.6 hr^-1^ with a media peristaltic pump (Buchler), while the volume in the culture was constantly removed at the same rate with a waste peristaltic pump (Zellweger Analytics). OD and mKO fluorescence readings were taken every 5 min for the duration of the experiment. Samples were taken periodically for CFU counting.

### Quantification of relative abundance of strains in continuous cultures

Standard growth curves measuring density (OD, absorbance at 650 nm) and fluorescence for WT and Δ*mexZ*-mKO strains were separately obtained in the continuous culture device without dilution (**Fig. S5A**). The plot of mKO fluorescence versus OD shows negligible fluorescence for the WT strain and a linear relationship between fluorescence and OD for the Δ*mexZ*-mKO strain within a window of ODs used in our experiments (**Fig. S5B**). The slope of this linear part of the plot was used to determine a conversion factor *S* between mKO fluorescence and the fraction of OD corresponding to the Δ*mexZ*-mKO strain, which allows us to calculate the relative abundance of Δ*mexZ* during competition with WT. We calculate the relative abundance of Δ*mexZ* in the culture 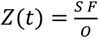, where *F* is the total fluorescence of the competition culture, *S* is the conversion factor, and *O* is the total OD of the competition culture.

### Response Dynamics assay

Glycerol stocks were inoculated into LB media and grown overnight, diluted 100-fold in 96-well microtiter plates (Falcon) containing 185 μl of fresh medium per well. OD (absorbance at 595 nm) was recorded by a plate reader (Synergy Neo-2) every 5 minutes for at least 36 hr. A concentration gradient of spectinomycin was set up over multiple 96-well plates. 15 μl of the spectinomycin gradient was added 30 min after OD was first detected above background. We used custom Matlab scripts to calculate the time required for the culture to double following a drug exposure, as well as the resulting steady-state growth rate of the population after growth was resumed, calculated by linear regression of log_2_(OD) during the remaining exponential growth. All relevant data was included in this study and made available online as supplemental data.

### Dynamic and Steady-state resistances

From the plate reader experiments, we extract two complementary measures of resistance: 1) *Dynamic Resistance*, which is the drug concentration that introduces a delay of one doubling time in the time to reach one doubling following drug exposure in comparison to the absence of drug, and 2) *Steady-State Resistance*, which is the drug concentration that halves the growth rate of the recovered population (similar to the IC_50_ concentration). We used custom Matlab scripts to fit the curves of delays and growth rates as functions of drug concentration, using exp1 and power2 respectively. These fits were then used to calculate dynamic and steady-state resistance. All relevant code is available online.

### Panel of representative *P. aeruginosa* strains

We imported a set of 47 *P. aeruginosa* specimens reflecting the organism’s diversity, obtained from the International Pseudomonas Consortium Database (IPCD). This set is mostly based on a panel that has been fully sequenced and previously characterized, which includes reference genomes, transmissible strains, sequential CF isolates, strains with specific virulence characteristics, and strains that represent serotypes, genotypes, or geographic diversity^12,43–45^. We added lab strains PA14 and PAO1 to the collection for a total of 49 strains. We used blast to retrieve the *mexZ* sequences from each of these genomes, which were then aligned using a custom Matlab script. We calculated alignment scores of each *mexZ* sequence to the consensus sequence using the Needleman-Wunsch algorithm. We classified strains as having a *mexZ* truncation if a truncation or large indel was observed (**Table 1**). We then ran our Response Dynamics assay in each strain as described above, using a spectinomycin gradient of 6.8 to 2143 μg/ml. 5 strains failed to grow in our culture medium formulation and were excluded from further analysis. Dynamic and steady-state resistances were calculated as described above, with projections beyond our drug range capped at 2500 μg/ml.

### Construction of the phylogenetic tree of *P. aeruginosa* strains

We determined the phylogenetic structure of our *P. aeruginosa* panel using 7 housekeeping genes *acsA, aroE, guaA, mutL, nuoD, ppsA, and trpE*, which are typically used to determine MLST types^67^. Gene sequences were retrieved from the sequenced genomes using blast and compiled into a single sequence. Pairwise distances were calculated using the Jukes-Cantor method, and the phylogenetic tree was inferred using an unweighted average distance method (UPGMA). Trees were plotted using iTOL.

### Phylogenetic independent contrasts (PIC)

We associated the presence of *mexZ* truncations with resistance phenotypes by performing trait comparative analyses^46,47^ using phylogenetic independent contrasts (PIC)^48^, which account for the underlying phylogenetic structure of the *P. aeruginosa* collection. We computed orthonormal contrasts^68^ for each trait using the library Ape in R. We then performed linear regression to establish correlations between the traits using custom Matlab script.

## Data Availability

The data that support the findings of this study are openly available. All data and code generated in the study are included in the SI and at our GitHub page: https://github.com/schultz-lab/Response-Dynamics. Further inquiries can be directed to the corresponding authors.

## Author Contributions

D.S. and D.R. designed the study. D.R. performed the microfluidics and continuous culture experiments. D.S. and D.R. analyzed the data. Y.D. created the knockout and reporter plasmids, and D.R. constructed the strains. R.M, M.B, J.V., and D.R. analyzed the IPCD collection of strains. D.S. and D.R. wrote the manuscript with input from all authors.

## Acknowledgements

We thank George O’Toole, Robert Cramer, and Chen Liao for the careful reading and valuable suggestions to our manuscript. We also thank Chris Geiger and Fabrice Jean-Pierre for important discussions on building our strains and experimental procedures, and Zdenek Svindrych for help with imaging. We thank Carey Nadell and George O’Toole for gifting us our ancestor strains. We thank Roger Lévesque for providing us the strains from the IPCD database.

