## Supplemental Information for "Heterogeneity during induction of resistance delays *Pseudomonas aeruginosa* recovery from antibiotic exposures"

### Common regulatory mutation increases single-cell survival to antibiotic exposures in *Pseudomonas aeruginosa*.

##### Fabrication of the microfluidic device

The device was fabricated by aligning a layer of 25 μm height containing the flow channels and a layer of 1.5 μm height containing the single-cell traps. The master mold was fabricated by ultraviolet photolithography using standard methods. SU-8 (Microchem) photoresist was applied to a silicon wafer by spin coating to appropriate thickness corresponding to the channel height, and patterns were then created by exposing the uncured photoresist to ultraviolet light through custom masks. We used a quartz mask for finer features (Toppan) and a film mask for the flow channels (CAD/Art Services). PDMS monomer (Sylgard 184) with 1:10 curing agent was mixed and poured on top of the wafer mold and air bubbles were removed using a vacuum chamber. After baking at 65°C for 3 hours, the PDMS was peeled off and inlet holes were introduced using a biopsy punch to connect the flow channels to the external tubing. Individual chips were then cut and bonded onto KOH-cleaned 1.5 coverslip slides (VWR) using oxygen plasma treatment, followed by 30 min on an 80°C hotplate and post-baked at 65°C overnight. Media was pumped through the device using a custom-built system that applies air pressure to media containers to induce flow through the device. We used an Arduino to control airflow through solenoid valves, allowing us to switch the pressures applied to containers of media with and without drug. The media inlets both feature high flow resistance channels to prevent backflow and have a short length of contact to prevent mixing (**Supplemental Video S1**).

##### Fabrication of the continuous culture device

The main body of the device is a custom-built 3D printed (Stratasys J55) housing, which has a central section to hold a glass vial containing the culture and a 4-pronged magnetic stirbar (IKA Works), as well as slots for LEDs and filter sets to measure two fluorescence channels, a 650 nm laser (Quarton) to measure OD, 3 photodiodes (TT Electronics) to quantify the fluorescence and OD readings, and a 3-D printed base stand underneath the main body, which houses a motor attached to magnetic beads to stir the culture. mKO fluorescence was quantified using a 525-50 nm emission filter (Chroma), a 600-50 nm excitation filter (Chroma), and 530 nm LED light source (LUXEON Star). We performed the experiments on a PA14 strain that has 2 chromosomal insertions of mKO (Fig. S4D). To amplify the photodiode signals, we used an inverting amplification circuit for OD quantification and an inverting amplification followed by a non-inverting amplification circuit for the fluorescence readings. A Raspberry Pi with a data acquisition board (Pi-Plate) was used to acquire fluorescence and OD measurements and to control the culturing system, using custom-built software.

##### Construction of plasmids and knockout strains

A *mexXY* reporter plasmid construct (P*mexXY*-YFP-P*rpoD*-CFP) was ordered from Genewiz and used as a base for our reporter system. Our reporter plasmid (DA350) uses a superfolder green fluorescent protein gene (sfGFP)[1], positioned downstream of the P*rpoD* to serve as a reference signal, and a red fluorescent protein gene (mCherry)[2] fused downstream of the *mexXY* promoter to measure its activity. The sfGFP gene was amplified via PCR from plasmid DA313[3] using primers P-GFP-F and P-GFP-R, while the mCherry gene was similarly amplified from plasmid JF72[4] using primers P-RFP-F and P-RFP-R. The backbone fragments were PCR amplified by primers sets P-bk-F1/P-bk-R1, P-bk-F2/P-bk-R2, and P-bk-F3/P-bk-R3, respectively, using P*mexXY*-YFP-P*rpoD*-CFP as template. All PCR fragments were purified and assembled into plasmids using the Gibson assembly method. Following transformation into competent *E. coli* NEB10beta cells (NEB), the resultant transformants were selected for ampicillin resistance by plating onto LB agar plates supplemented with 100 µg/mL of ampicillin. The accuracy of all plasmid sequences was confirmed by Sanger DNA sequencing. This new reporter plasmid offered improved resolution over background fluorescence in *P. aeruginosa*, enabling more accurate measurement of P*mexXY* promoter activity.

The plasmids used to knock out *mexZ* and *mexY* genes from *P. aeruginosa* PA14 strain were constructed using pMQ30 as backbone[5]. The ~1kb upstream and downstream fragments of the target gene were PCR amplified from genomic DNA of PA14 and Gibson assembled into the backbone of pMQ30. Plasmid DA354 was created to knock out *mexZ*. Specifically, two backbone fragments were amplified via PCR from PMQ30 using primer sets Mq-bk-F1/Mq-bk-R1 and Mq-bk-F2/Mq-bk-R2. The upstream and downstream DNA regions of *mexZ* were PCR amplified with primer sets Z-up-F/Z-up-R and Z-dw-F/Z-dw-R, respectively. The four fragments were assembled to make the resultant plasmid DA354. Similarly, plasmid DA355 was constructed by fusing the upstream and downstream DNA regions of *mexY*, together with the same pMQ30 backbone. The upstream and downstream regions of *mexY* were PCR amplified with primer sets Y-up-F/Y-up-R and Y-dw-F/Y-dw-R, respectively. The constructs were then transformed into *E. coli* NEB10beta cells (NEB) and spread onto LB plates with gentamicin (10 µg/mL). Following a 24-hour incubation period at 30°C, resistant colonies were selected and propagated in LB broth supplemented with the same antibiotic at 30°C, facilitating plasmid extraction. Subsequently, the purified plasmids were verified through Sanger sequencing to ensure accuracy.

The Δ*mexZ* and Δ*mexY* knockout strains were generated through in-frame deletions to avoid any potential downstream polar effects, using the above plasmids DA354 and DA355, respectively. The gene knockout followed a previous published two-step allelic exchange protocol[6]. Briefly, the plasmid was first transformed to *E. coli* S17 strain, after which it was conjugated with *P. aeruginosa* PA14 and selected for gentamicin resistance at 30 µg/mL. The knockout out mutants were then subjected to counter selection using sucrose. The right knockout mutants were confirmed by colony PCR[6].

All plasmids were constructed by Gibson assembly[7] using NEBuilder® HiFi DNA Assembly kit (NEB). PCR reactions were conducted using high-fidelity Q5 DNA polymerase (NEB) and the products were purified prior to Gibson assembly. All PCR primers used in this study are listed in Supplemental Table S1. When necessary, primers were specifically designed to include overlapped regions for the assembly process. The assembled plasmids were transformed into chemically competent *E. coli* NEB10beta cells (NEB) and selected for appropriate antibiotic resistances. Plasmids, PCR products, and DNA fragments from agarose gel were purified with Qiagen miniprep, PCR purification, and Gel extraction kits, respectively.

| Primers | Sequences 5’-----3’ |
|---|---|
| P-GFP-F | tttgtaaagtagacctaaggagtaaatatgagcaaaggagaagaacttttcactgg |
| P-GFP-R | ccgatcaagtcttcgcgatgattattatttgtagagctcatccatgccatgtgt |
| P-RFP-F | acccccggtgcagaaaataaggaggaaaaaaaaatggtgagcaagggcgaaga |
| P-RFP-R | gacctgcattacttgtacagctcgtccatgccgc |
| P-bk-F1 | gcggcatggacgagctgtacaagtaatgcaggtcgtctcggatcgagaagg |
| P-bk-F2 | ggtgatgacggtgaaaacctctga |
| P-bk-F3 | gctcatatttactccttaggtctactttacaaaaaataagcagaggattatacctgatg |
| P-Bk-R1 | tcagaggttttcaccgtcatcaccgaaacgc |
| P-bk-R2 | ataataatcatcgcgaagacttgatcggtgccgggat |
| P-bk-R3 | tattttctgcaccgggggtgtccctcgattc |
| Y-up-F | tcagaccgcttctgcgttctgacccttgaaggcggccctgga |
| Y-up-R | agcaccggcggcgcgatgtcc |
| Y-dw-F | acatcgcgccgccggtgctggtaccgctgttcttcctgg |
| Y-dw-R | tgcgttggccgattcattcaggacggcgaattgctgcgac |
| Mq-bk-F1 | gttccgcgcacatttccccgaaacctctgacacatgcagctcccgg |
| Mq-bk-F2 | ctgaatgaatcggccaacgcacgggaagag |
| Mq-bk-R | tcagaacgcagaagcggtctgat |
| Mq-bk-R2 | ttcggggaaatgtgcgcggaacc |
| Z-up-F | gttttatcagaccgcttctgcgttctgagcacgcatggcgatggtg |
| Z-up-R | aggtagggagaactgcgcaggctgtcgcgggttttctgggattc |
| Z-dw-F | agcctgcgcagttctccct |
| Z-dw-R | tcccgtgcgttggccgattcattcaggaacggcgcgtagagctgg |

**Supplemental Table 1 -** List of PCR primers used in this study.

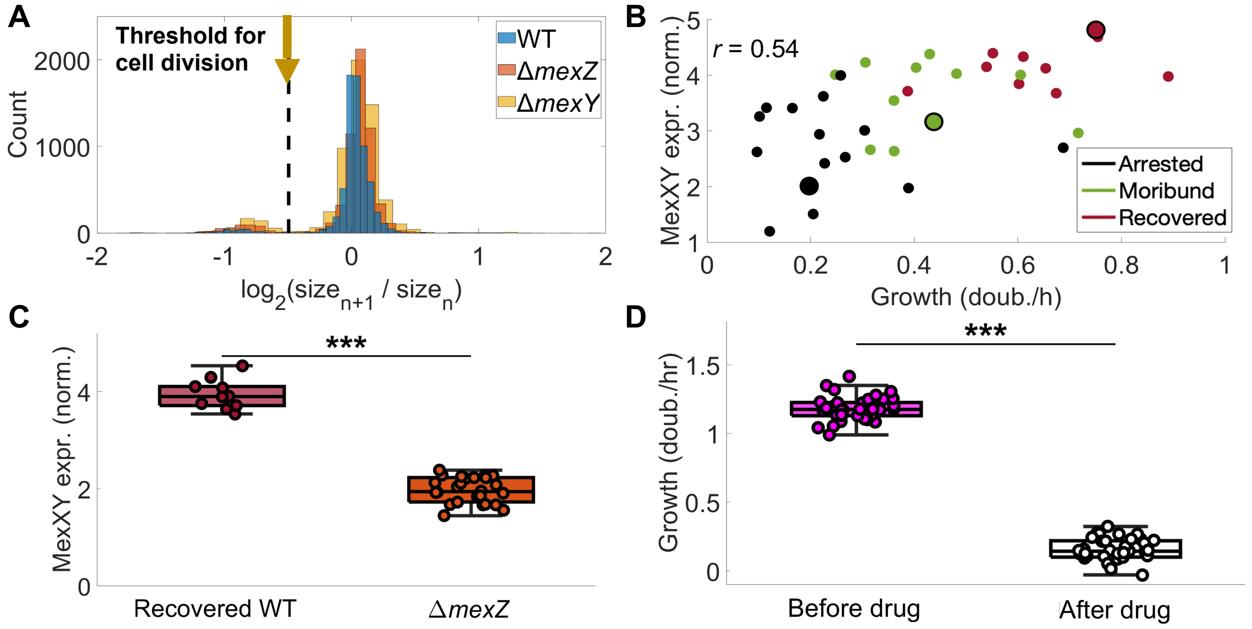


**Supplementary Figure 1 - Heterogeneous cell growth and gene expression in the microfluidic experiment. (A)** Histogram of cell sizes relative to their size at the previous time step (10 minutes earlier), throughout the whole experiment for all genotypes in the microfluidic assay. The dotted line depicts the threshold used to identify division events. **(B)** Maximum MexXY expression and final growth rate of all WT cells. The Recovered, Moribund, and Arrested cells depicted in **Fig 1C** in the main text are denoted by large dots. Pearson correlation between final levels of *mexXY* expression and growth rate is displayed. **(C)** Comparison of final MexXY expression of Recovered WT cells and Δ*mexZ* cells. **(D)** Comparison of all WT cells' growth rates before and after drug exposure. The asterisks denote *** $p < 0.001$ (**Methods**).

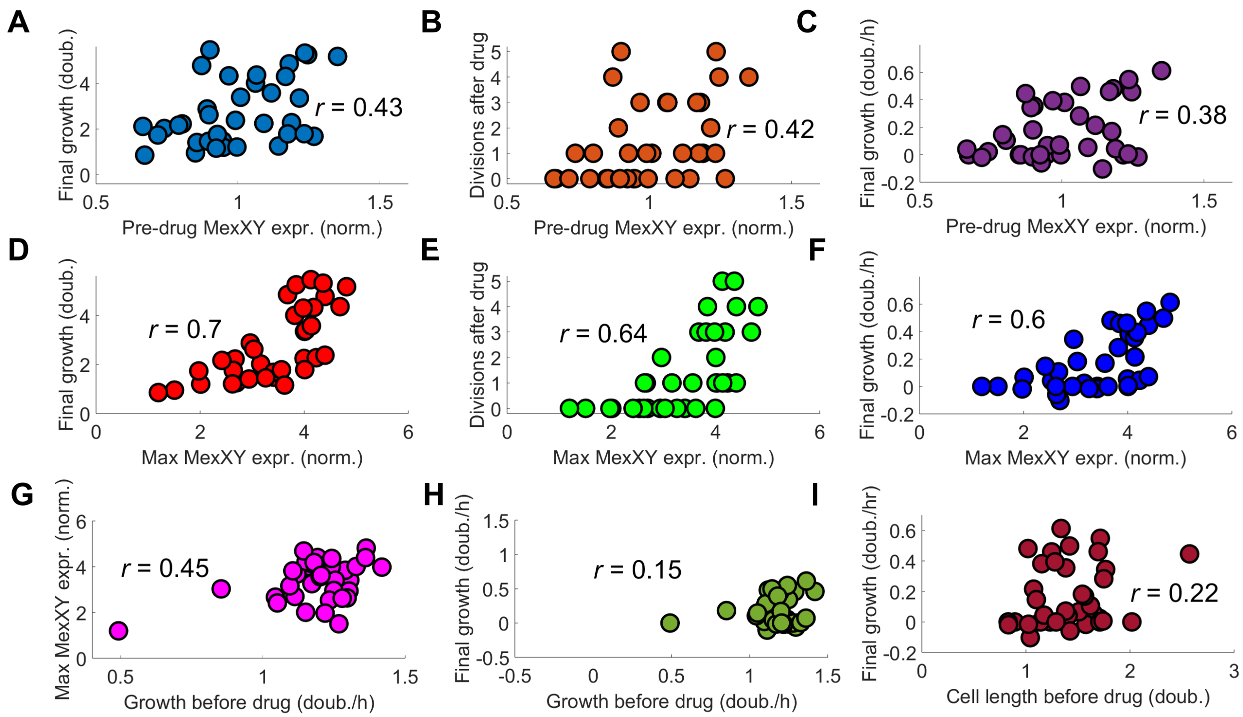


**Supplementary Figure 2 - Induction of MexXY expression promotes recovery of cell growth following antibiotic exposure.** In WT cells tracked in the microfluidic device, **(A)** accumulated growth, **(B)** division events, and **(C)** final growth rate have a moderate positive correlation with MexXY expression at the moment of drug exposure. However, **(D)** accumulated growth, **(E)** division events, and **(F)** final growth rate have a high positive correlation with the maximum MexXY level reached after drug exposure. **(G)** Maximum MexXY expression correlates moderately with the growth rate at the time of drug exposure, but **(H)** final growth rate does not correlate with the growth rate at the time of drug exposure. **(I)** The final growth rate also does not correlate with cell length at the moment of drug exposure. Pearson's correlation coefficient is displayed on each graph.

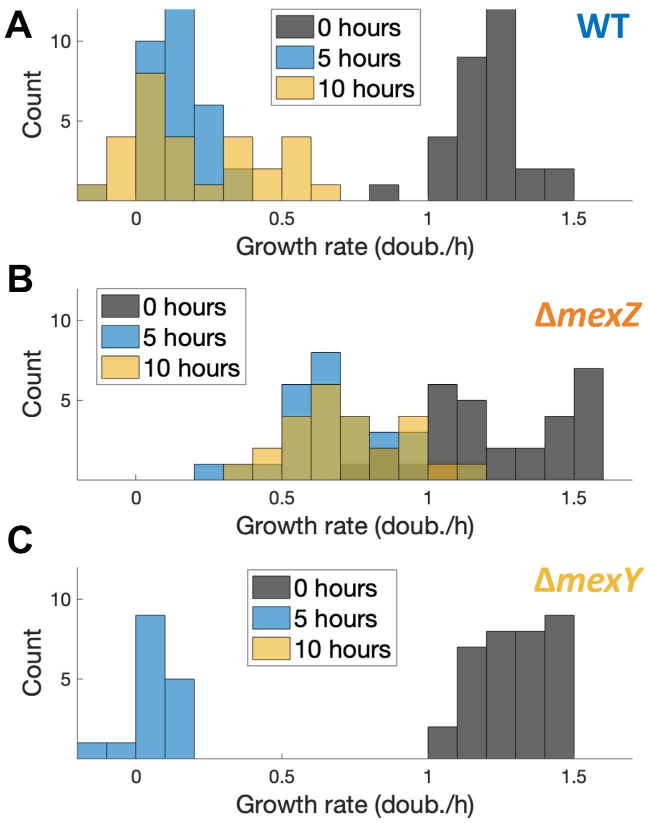


**Supplementary Figure 3 - *mexZ* deletion increases survival to an abrupt exposure to spectinomycin.** Histogram showing distribution of growth rates for **(A)** WT, **(B)** Δ*mexZ*, and **(C)** Δ*mexY* cells following exposure to 1000 μg/mL spectinomycin in the microfluidic assay. Histograms are shown at the time of exposure (0 hours), as well as 5 and 10 hours afterwards. WT cells display a significant reduction of growth followed by heterogeneous recovery, while Δ*mexZ* cells maintain growth throughout the experiment.

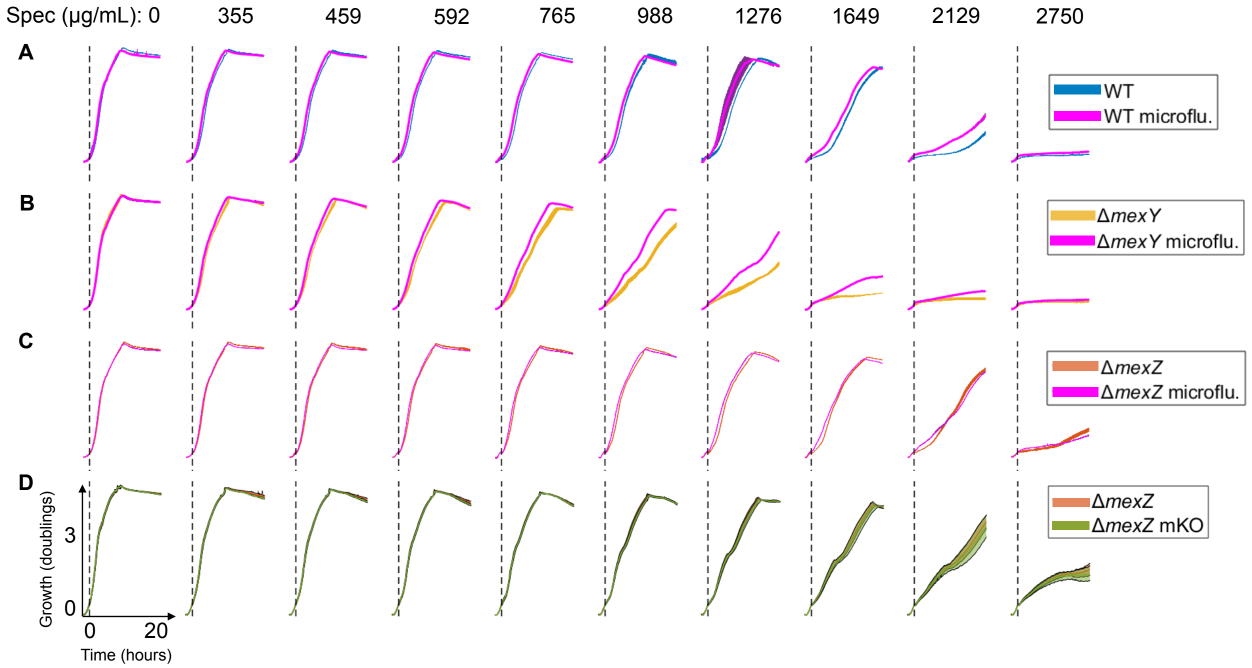


**Supplementary Figure 4 - Deletion of motility genes, reporter plasmids, and mKO chromosomal insertions do not affect *P. aeruginosa*'s growth or spectinomycin resistance.** Comparison of growth in **(A)** WT, **(B)** Δ*mexY*, and **(C)** Δ*mexZ* strains to their counterparts used in the microfluidic assays, which have a deletion of motility genes and contain a MexXY reporter plasmid. The dotted lines denote abrupt additions of spectinomycin at the doses specified above the panels. We did not detect significant differences in cell growth or resistance. **(D)** Similarly, the Δ*mexZ* strain used in the continuous cultures, which carries 2 chromosomal insertions of mKO, shows similar growth and resistance profile as the Δ*mexZ* strain from which it is derived. The shading around each line graph represents the maximum and minimum of two replicates.

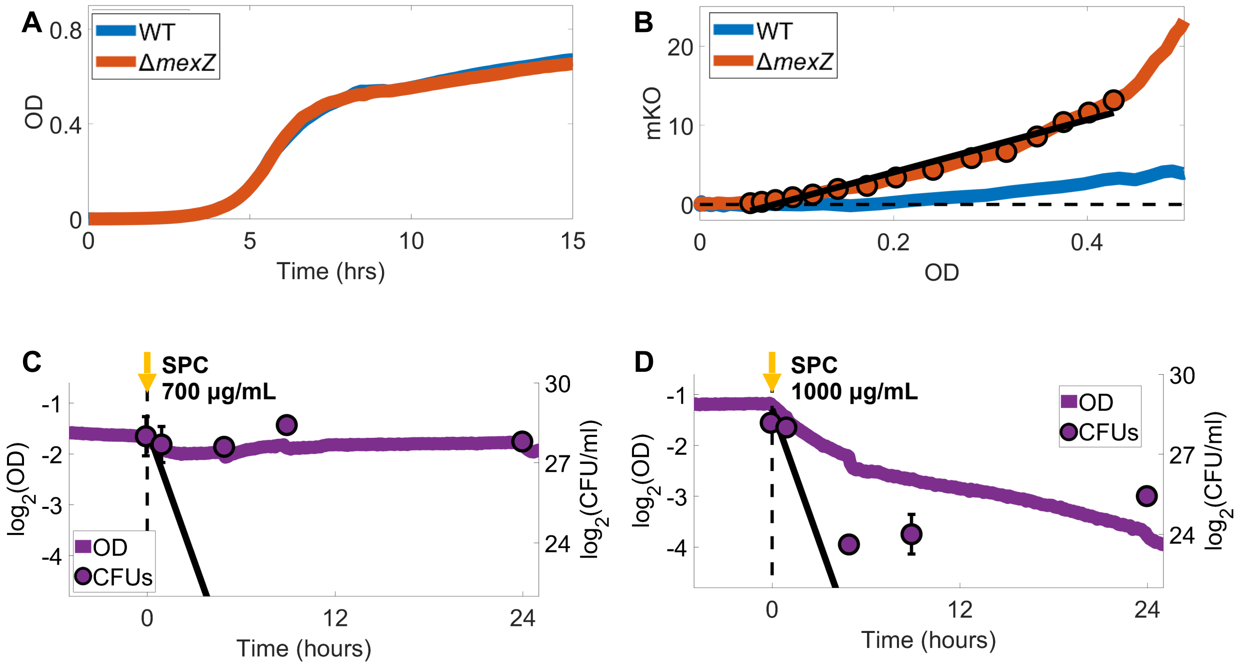


**Supplementary Figure 5 - Calibration of the continuous culturing device. (A)** Growth of monocultures of a WT strain and a Δ*mexZ* strain with two chromosomal insertions of mKO, in the absence of dilution. **(B)** Fluorescence measured from the Δ*mexZ* strain increases linearly with OD within the range used in our experiments. The slope of the black fitted line overlaid on Δ*mexZ*'s standard curve was used to determine Δ*mexZ*'s relative abundance during competition assays in Fig 3DE in the main text (**Methods**). Competition cultures were kept at 0.4 OD or below, minimizing the effect of WT auto-fluorescence on readings. **(C-D)** Total OD of the mixed culture during the competition assays between WT and Δ*mexZ* under **(C)** 700 µg/mL and **(D)** 1000 µg/mL spectinomycin, from Fig 3DE in the main text. Dotted line denotes the moment of drug exposure. Black line denotes the limit of dilution in the absence of growth.

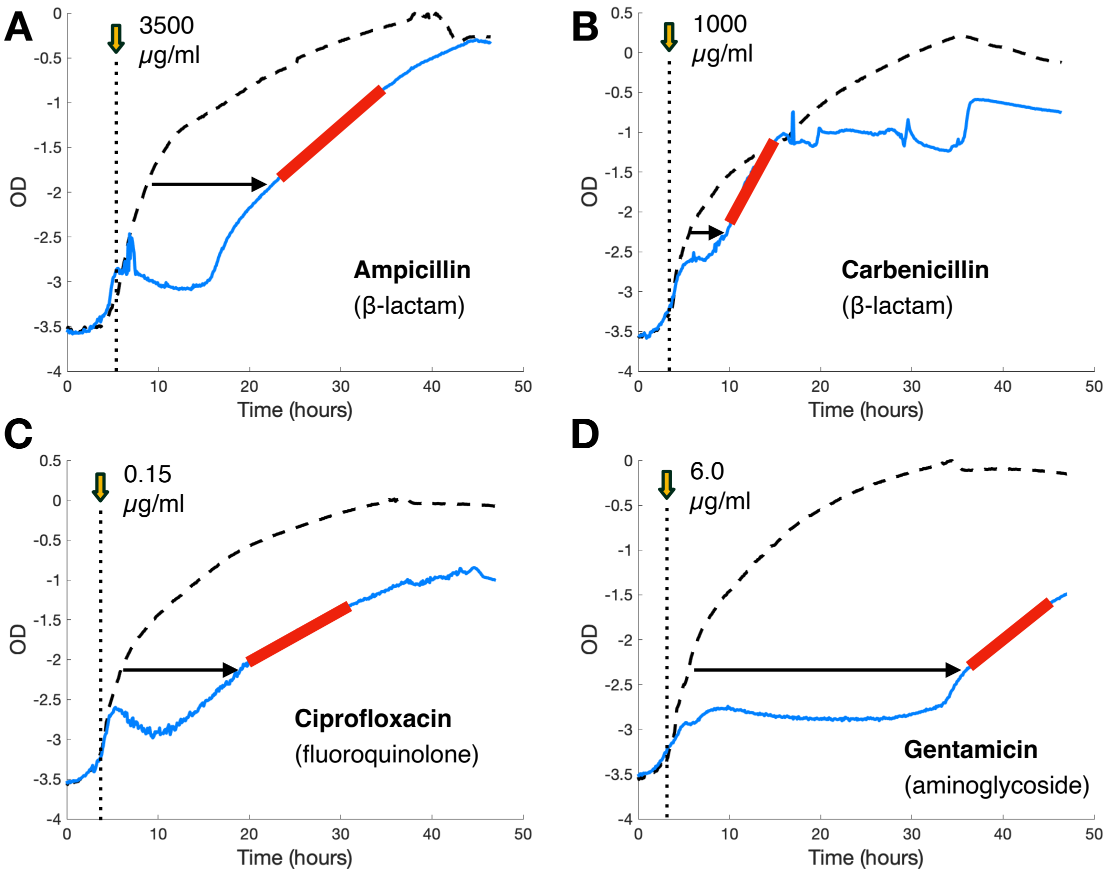


**Supplementary Figure 6 – Delay in population-level growth recovery is general across different strains and drug classes.** Growth curves of *P. aeruginosa* strain PA14 (WT) following exposures to **(A)** ampicillin (β-lactam), **(B)** carbenicillin (β-lactam), **(C)** ciprofloxacin (fluoroquinolone) and **(D)** gentamicin (aminoglycoside). The vertical dotted line shows the time of drug exposure. Fluoroquinolones and β-lactams are bactericidal drugs and often result in a reduction of OD following exposure.

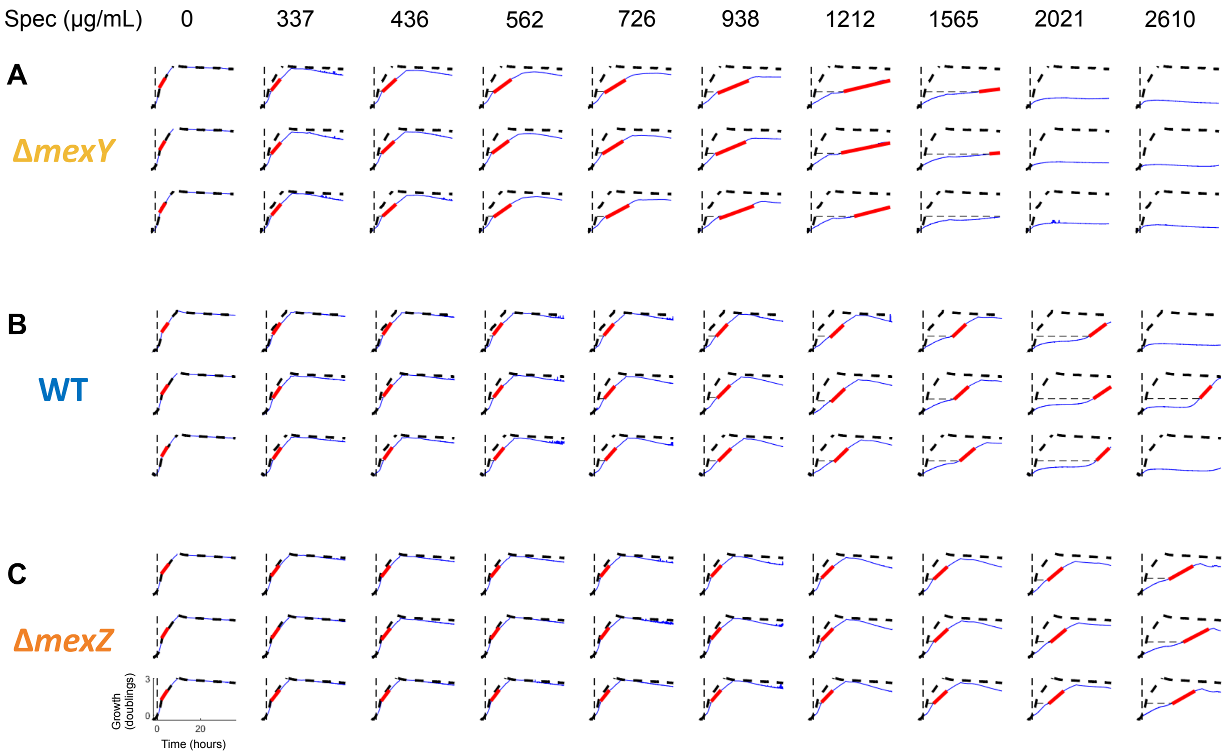


**Supplementary Figure 7 - Calculation of Steady-State and Dynamical resistances from cell growth during an abrupt drug exposure.** Growth curves of **(A)** Δ*mexY*, **(B)** WT, and **(C)** Δ*mexZ* cells following exposure to spectinomycin concentrations picked from a gradient, performed in three replicates. The red line shows the steady-state growth following population recovery, which was used to calculate Steady-state resistance. The horizontal dotted line shows the delay in growth recovery in comparison to the absence of drug, which was used to calculate Dynamical resistance. The overlaid black dotted line represents growth in the absence of drug, which is the first column of each row. These growth curves were used to calculate delays and steady-state growth in Fig 4B-D in the main text.

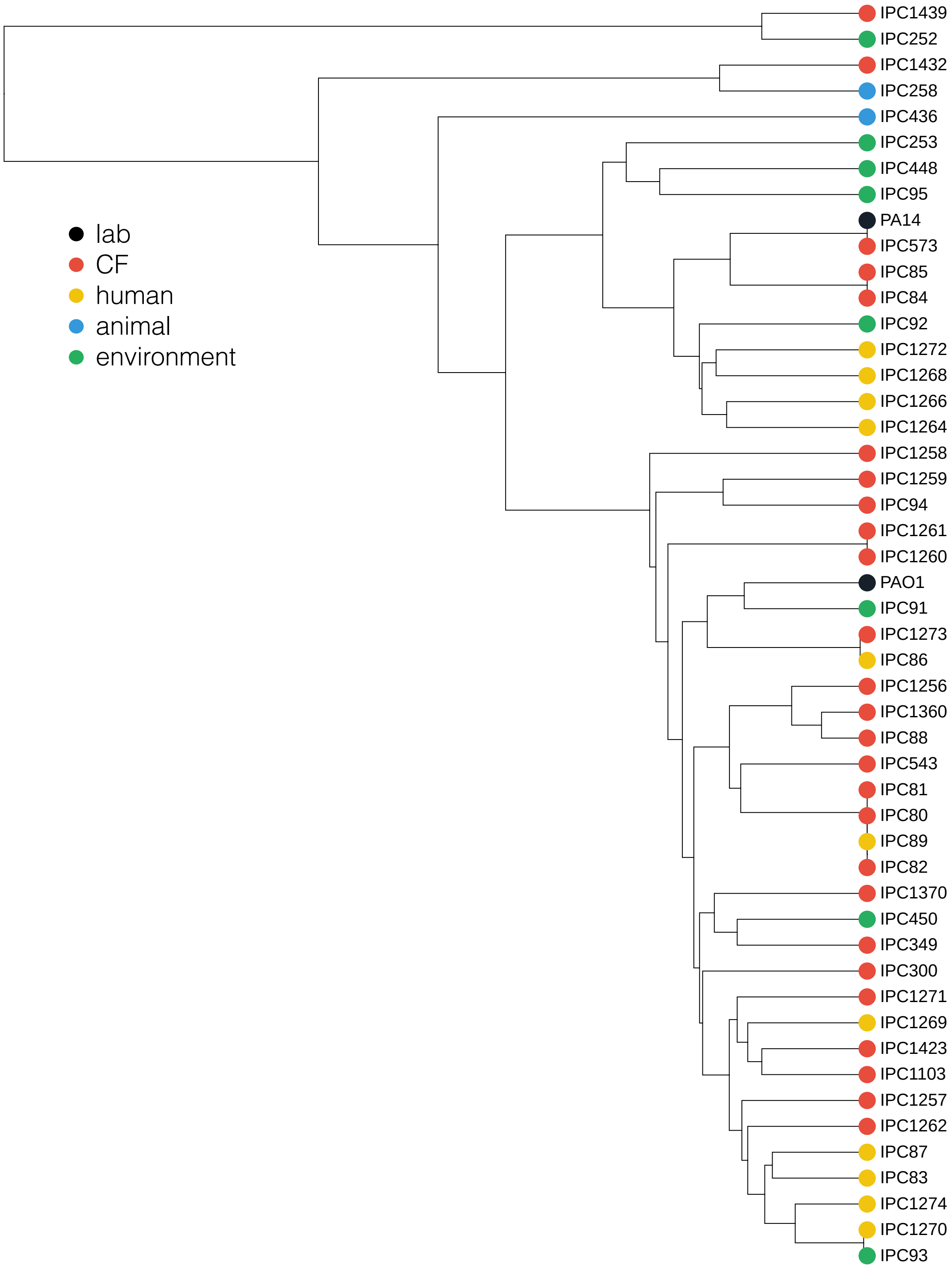


**Supplementary Figure 8 – Phylogenetic tree of *P. aeruginosa* specimens.** A set of 49 *P. aeruginosa* specimens reflecting the organism's diversity was obtained from the International Pseudomonas Consortium Database (IPCD). We determined the phylogenetic structure of our *P. aeruginosa* panel using 7 housekeeping genes typically used to determine MLST types. Inset shows environments where samples were obtained.

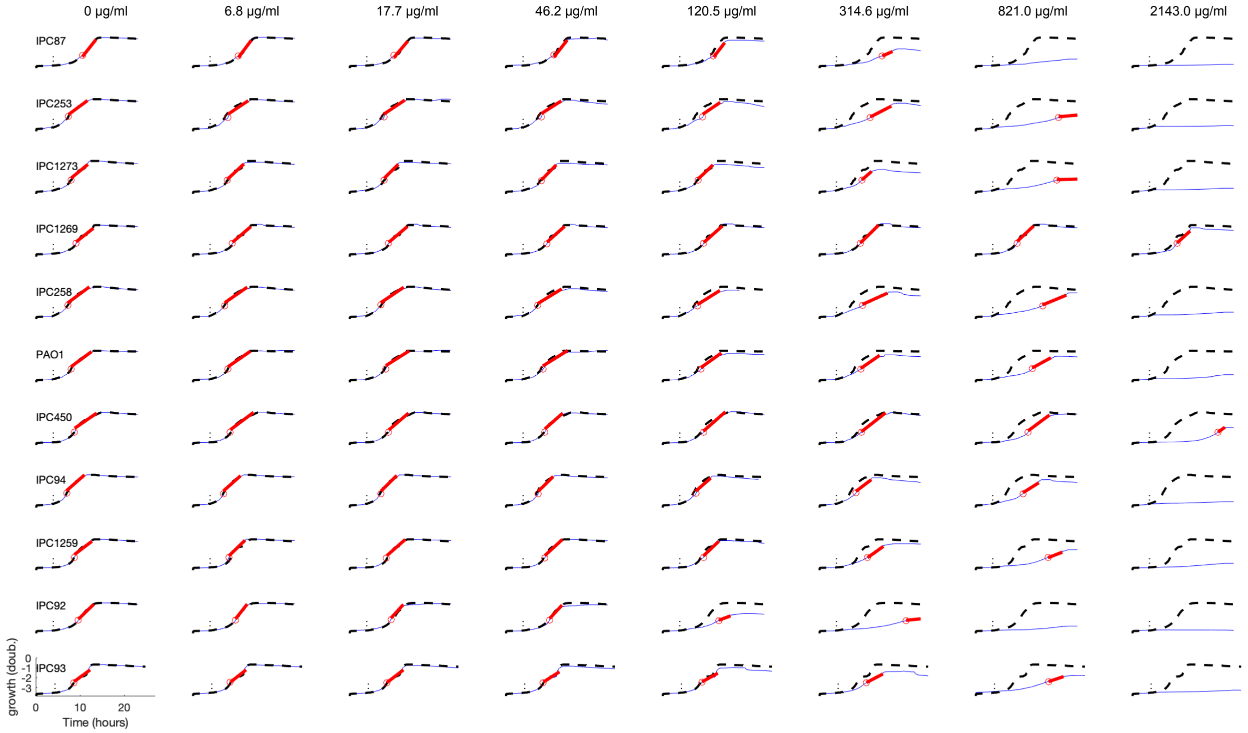


**Supplementary Figure 9 – Calculation of Steady-State and Dynamical resistances for strains in the *P. aeruginosa* panel – Plate 1.** Growth curves of isolates following exposure to spectinomycin concentrations picked from a gradient, indicated above. The red line shows the steady-state growth following population recovery, which was used to calculate Steady-state resistance. The horizontal dotted line shows the delay in growth recovery in comparison to the absence of drug, which was used to calculate Dynamical resistance. The overlaid black dotted line represents growth in the absence of drug, which is the first column of each row. These growth curves were used to calculate delays and steady-state growth in Fig. 5 in the main text.

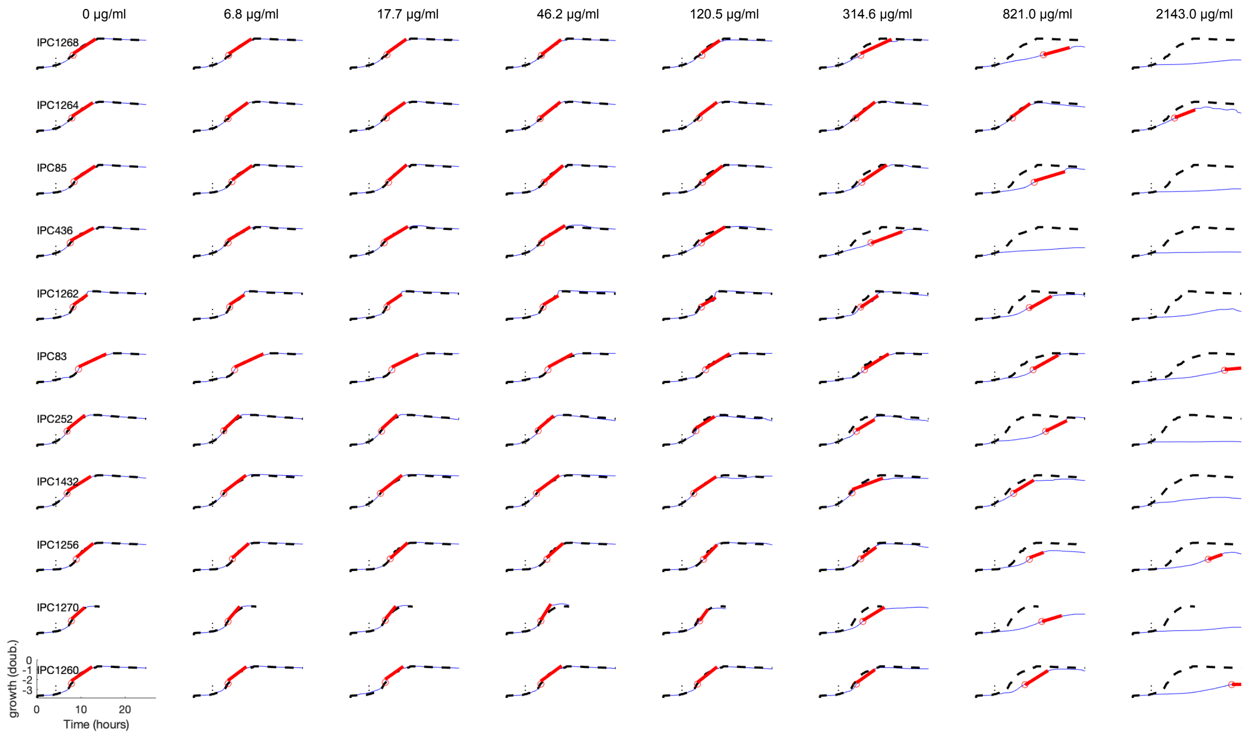


**Supplementary Figure 10 – Calculation of Steady-State and Dynamical resistances for strains in the *P. aeruginosa* panel – Plate 2.** Growth curves of isolates following exposure to spectinomycin concentrations picked from a gradient, indicated above. The red line shows the steady-state growth following population recovery, which was used to calculate Steady-state resistance. The horizontal dotted line shows the delay in growth recovery in comparison to the absence of drug, which was used to calculate Dynamical resistance. The overlaid black dotted line represents growth in the absence of drug, which is the first column of each row. These growth curves were used to calculate delays and steady-state growth in Fig. 5 in the main text.

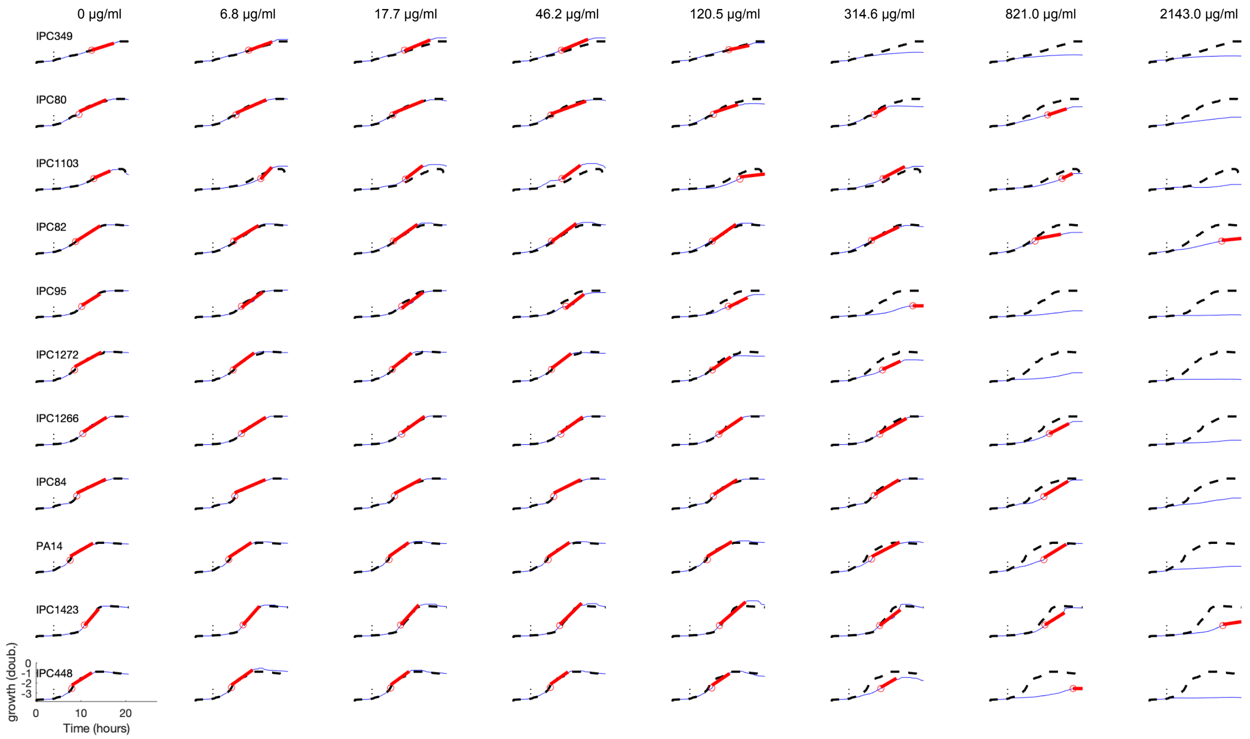


**Supplementary Figure 11 – Calculation of Steady-State and Dynamical resistances for strains in the *P. aeruginosa* panel – Plate 3.** Growth curves of isolates following exposure to spectinomycin concentrations picked from a gradient, indicated above. The red line shows the steady-state growth following population recovery, which was used to calculate Steady-state resistance. The horizontal dotted line shows the delay in growth recovery in comparison to the absence of drug, which was used to calculate Dynamical resistance. The overlaid black dotted line represents growth in the absence of drug, which is the first column of each row. These growth curves were used to calculate delays and steady-state growth in Fig. 5 in the main text.

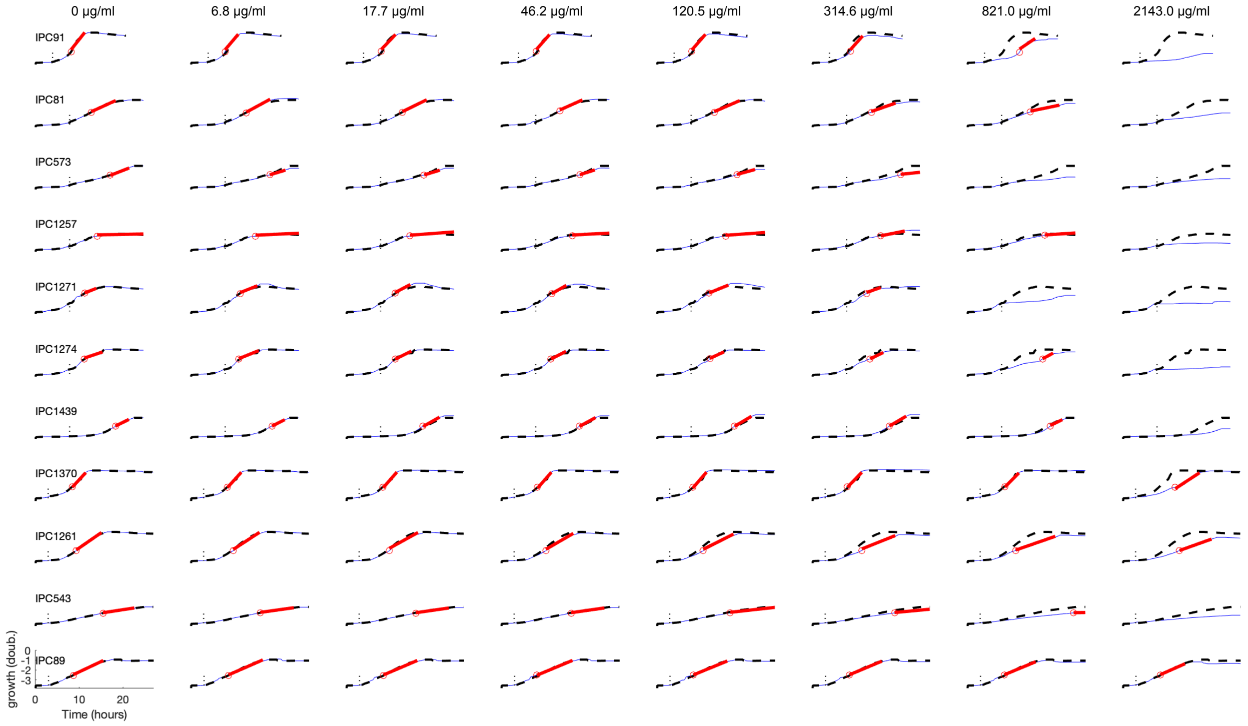


**Supplementary Figure 12 – Calculation of Steady-State and Dynamical resistances for strains in the *P. aeruginosa* panel – Plate 4.** Growth curves of isolates following exposure to spectinomycin concentrations picked from a gradient, indicated above. The red line shows the steady-state growth following population recovery, which was used to calculate Steady-state resistance. The horizontal dotted line shows the delay in growth recovery in comparison to the absence of drug, which was used to calculate Dynamical resistance. The overlaid black dotted line represents growth in the absence of drug, which is the first column of each row. These growth curves were used to calculate delays and steady-state growth in Fig. 5 in the main text.

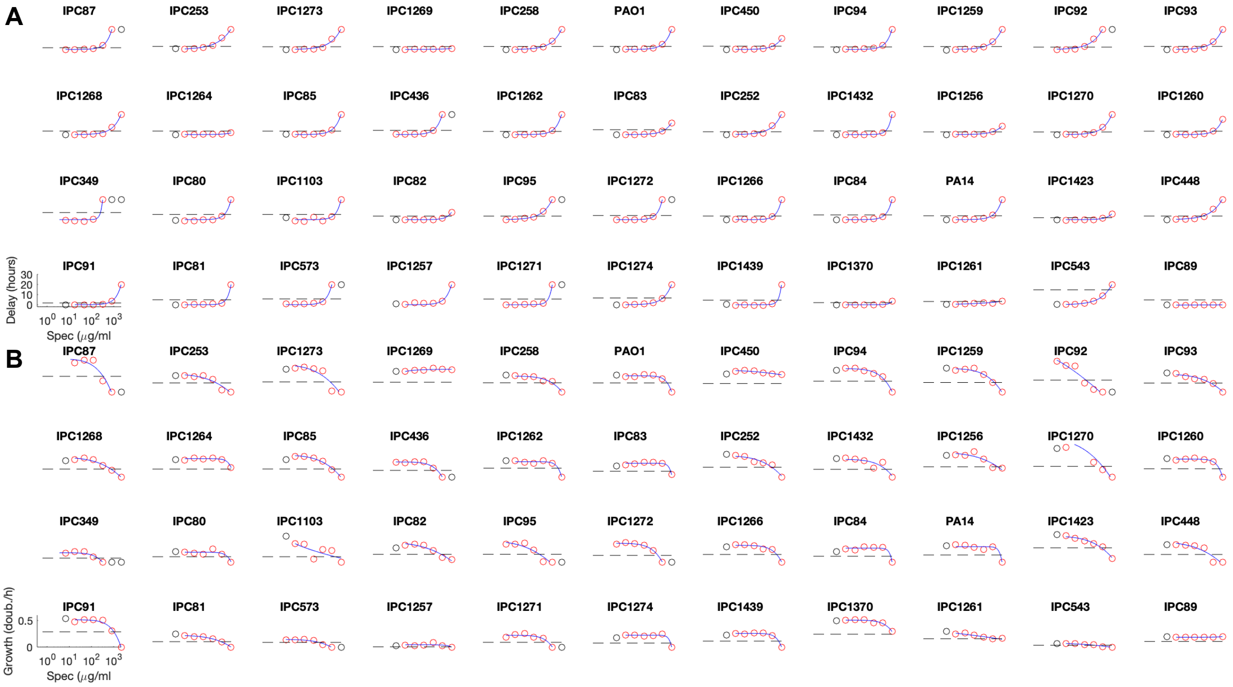

**Supplementary Figure 13 – Calculation of Steady-State and Dynamical resistances for strains in the *P. aeruginosa* panel. (A)** Recovery delays of each strain as a function of the spectinomycin dose used in the exposure. The horizontal line indicates the threshold of one doubling time, measured during exponential growth in the absence of drug. We calculate *Dynamical Resistance* as the drug concentration that causes a delay in recovery equal to this threshold. **(B)** Steady-state growth of each strain in our panel as a function of the spectinomycin dose used in the exposure. The horizontal line indicates the threshold of half of the maximum growth rate, measured during exponential growth in the absence of drug. We calculate *Steady-state Resistance* as the drug concentration that reduces the steady-state growth rate to this threshold.

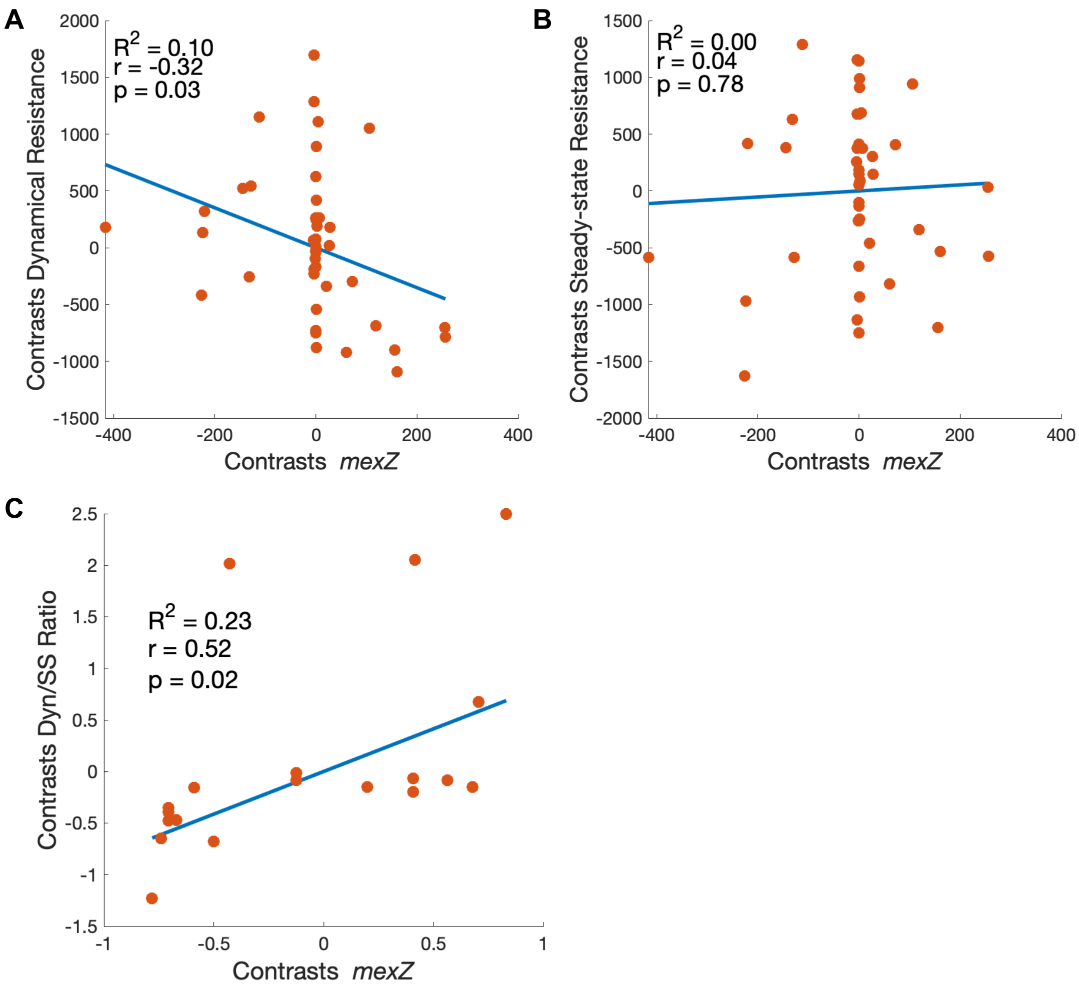


**Supplementary Figure 14 – *mexZ* truncations are associated with increased dynamical resistance.** **(A)** We calculated phylogenetic independent contrasts (PIC) of the presence of *mexZ* variants using the alignment score of each variant to the consensus sequence, which is a continuous trait. Dynamical resistance still correlates with the presence of *mexZ* variants more distant from the consensus sequence (lower alignment scores), but no extra information is gained in comparison to contrasts calculated using the presence of *mexZ* truncations, a binary trait. **(B)** No correlation is found between steady-state resistance and the presence of distant *mexZ* variants. **(C)** Correlation between the contrasts of the ratio of dynamical and steady resistance and the binary presence of *mexZ* truncations.

**Supplementary Video 1- Microfluidic device allows rapid switching between media.** In the slowed video, the top media stream is composed of water (no color) and the bottom media stream is composed of water with fluorescein (green color). Both fluid streams are laminar and meet for a short distance, causing minimal mixing. By switching the pressure applied to each media container, we can rapidly select which medium reaches the channel feeding the single-cell traps.

**Supplementary Video 2 - Diversity of fates in WT single cells responding to a step increase in spectinomycin concentration.** WT cells carrying the native *mexXY* resistance mechanism were exposed to a step increase of 1000 µg/mL spectinomycin at time zero. A fluorescent reporter with a matching promoter was used to measure expression of the efflux pump component MexY (red, mCherry). Imaging was done in a microfluidic device that places single cells in fixed locations as they go through division cycles. Valves allow on-chip rapid switching of media.

**Supplementary Video 3 - Δ*mexZ* cells survive a step increase in spectinomycin concentration.** Cells carrying a Δ*mexZ* deletion were exposed to a step increase of 1000 µg/mL spectinomycin at time zero. Unlike WT cells, all Δ*mexZ* cells survived the exposure and maintained growth under the drug.

**Supplementary Video 4 - Δ*mexY* cells do not survive a step increase in spectinomycin concentration.** Cells carrying a Δ*mexY* deletion were exposed to a step increase of 1000 µg/mL spectinomycin at time zero. Without the resistance provided by MexXY, all Δ*mexY* cells quickly stopped growing following drug exposure.

**Supplementary Video 5 - WT cells exposed to 700 µg/mL spectinomycin show reduced growth.** WT cells carrying the native *mexXY* resistance mechanism and a MexY reporter plasmid were exposed to a step increase of 700 µg/mL spectinomycin at time zero. WT cells mostly show reduced growth but survive this drug concentration, unlike the heterogeneous cell fates of WT cells exposed to 1000 µg/mL spectinomycin.
